# Recombination, selection and the evolution of tandem gene arrays

**DOI:** 10.1101/2022.01.26.477888

**Authors:** Moritz Otto, Yichen Zheng, Thomas Wiehe

## Abstract

Multi-gene families – immunity genes or sensory receptors, for instance – are often subject to diversifying selection. Allelic diversity may be favoured not only through balancing or frequency dependent selection at individual loci, but also by associating different alleles in multi copy gene families. Using a combination of analytical calculations and simulations, we explored a population genetic model of epistatic selection and unequal recombination, where a trade-off exists between the benefit of allelic diversity and the cost of copy abundance. Starting from the neutral case, where we showed that gene copy number is Gamma-distributed at equilibrium, we derived also mean and shape of the limiting distribution under selection. Considering a more general model which includes variable population size and population substructure, we explored by simulations mean fitness and some summary statistics of the copy number distribution. We determined the relative effects of selection, recombination and demographic parameters in maintaining allelic diversity and shaping mean fitness of a population. One way to control the variance of copy number is by lowering the rate of unequal recombination. Indeed, when encoding recombination by a rate modifier locus, we observe exactly this prediction. Finally, we analyzed the empirical copy number distribution of three genes in human and estimated recombination and selection parameters of our model.

## Introduction

Multi-gene families occur in most, if not all, genomes of eukaryotes – in metazoans as well as in plants. They may be conserved across large evolutionary distances, such as the histones or tRNA gene families, or rapidly diversify in single species, such as the NLR-genes in *Danio rerio* (Howe *et al*. 2016) or the LRR-genes in *Arabidopsis thaliana* (de Weyer *et al*. 2019).

Interspecies comparison of gene families derived from whole genome duplication has been used, for instance, to estimate relative rates of gene loss and functional divegence (Nadeau and Sankoff 1997). On a shorter time scale, segmental duplication and unequal recombination are perhaps the more important mechanisms to explain gene family size differences between species, populations and individuals. Modeling gene family evolution has a quite long history (Smith 1974; Innan 2009; Demuth and Hahn 2009; Liu *et al*. 2011). The roadmap in a population genetic framework was laid out in a series of contributions by Ohta (Ohta 1976, 1979, 1984, 1987, 1988, 2000). These models typically include forces such as selection and unequal recombination or gene conversion. To describe the dynamics of copy number variation (CNV) generated by unequal recombination Takahata (Takahata 1981) introduced a general model based on the work of Krüger and Vogel (Krüger and Vogel 1975). Fostered especially by the human genome diversity projects, leading to the realization that structural variation is more than abundant in human populations and observing genome size differences between individuals of 100Mb and more (Tuzun *et al*. 2005; Redon *et al*. 2006; Eichler 2008), we are witnessing revived interest in modeling and analyzing the evolution of gene families and of the forces and mechanisms driving copy number polymorphisms.

Tandem gene duplication may happen due to some form of replication error, mis-pairing or segregation anomaly, notably unequal or – less frequently – non-homologous recombination (Silver 2001). A duplicated gene initially arises in a single individual, very much like a base mutation, and may be lost by drift or be propagated to the offspring in subsequent generations. On its way to fixation, or loss, such a duplication manifests itself as copy number variation (CNV) in a given population and – given sufficiently large populations – is sensed by the filter of natural selection. When beneficial, directional selection will accelerate its fixation and subsequent purifying selection will prevent it from loss. Alternatively, when beneficial only in conjunction with other alleles or other copies, balancing selection may force it to remain at intermediate frequency. The best known examples are perhaps the alleles of pathogen receptors and immune genes, such as those of the MHC complex in vertebrates. Balancing selection, however, comes with a fitness cost in terms of segregation load. Haldane had suggested (Haldane 1937) that this effect may be alleviated or avoided when overdominant alleles are arrayed in tandem on the same chromosome rather than be combined on homologous chromosomes. Only recently, this fundamental idea has been experimentally tested – and confirmed – in populations of the mosquito *Culex pipiens* (Milesi *et al*. 2017).

Here, we designed a model of tandemly arrayed genes whose evolution is driven by unequal recombination together with a mixture of diversifying and negative selection. More precisely, negative selection will keep copy number in check, while allelic diversity is positively selected. We implement this via a product of two multiplicative fitness components: one of them is decreasing with copy number and the other one is increasing with allele number (see eq. (1)). In its structure this fitness function is an old acquaintant. Very similar versions feature in the classical model of Muller’s ratchet (Haigh 1978) and its epistatic relatives (Kondrashov 1982; Chao 1988).

We discovered the following: first, in the absence of selection, i.e. when diversity of alleles does not confer any fitness benefit and additional copies do not provide any cost, the distribution of copy numbers can be analytically expressed. It is a Gamma distribution with shape *α* = 4 and with a scale which depends only on the mean copy number of the initial distribution. With selection, the limiting distribution is still well approximated by a Gamma distribution, but depends on the combination of selection coefficients and recombination rate, and *not* on the initial distribution. Second, population size can have a stronger effect on mean fitness and allelic diversity than the strength of selection itself. Third, low recombination rates may be favourable to maintain allelic diversity. Consistent with this, when recombination rates are coded as alleles at a modifier locus and are allowed to evolve over time, we observe a tendency towards recombination rate reduction.

Taken together, our model captures essential aspects of a multi gene family driven by a force of increasing allelic diversity and, at the same time, an opposing force of maintaining genome and chromosome integrity and of limiting both segregation and recombination loads.

Based on the empirical copy number distribution in a set of three exemplary gene families in human we estimated the strengths of selection and (unequal) recombination rates in a natural population.

## Methods

### Model

We consider a *compound* model in which the number of copies (*y*) of a certain gene per individual, as well as the number of alleles (*x*), are variable. When alleles are all considered distinct (but without labeling their identities) and copy numbers remain variable, we call this the *y-only model*.

In a diploid population of effective size *N* ≤ ∞ let individual *i*, 1 ≤ *i* ≤ *N*, carry 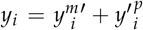 copies of a particular gene on its maternal (*m*) and paternal (*p*) chromosomes. We use the notation *y*′ for the number of copies per chromosome when neither the individual nor the parental status matter. If copies are distinguishable, we call them *alleles* and let *x*′, 1 ≤ *x*′ ≤ *y*′, be the number of different alleles on a chromosome with *y*′ copies. By extension, individual *i* has 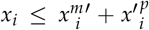 alleles (Fig 1C, alleles indicated by different colours). Fitness *ω*_*i*_ of individual *i* is determined by both copy and allele numbers: *ω*_*i*_ = *ω*_*i*_ (*x*_*i*_, *y*_*i*_). We assume that increasing the number of copies incurs a fitness *cost*, representing adverse effects to genomic structure and integrity, while increasing the number of alleles incurs a fitness *benefit*, representing improved function such as recognition of a wider range of pathogens or stimuli. To fix ideas, we consider the following fitness function

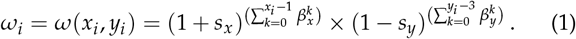

**Figure 1.**
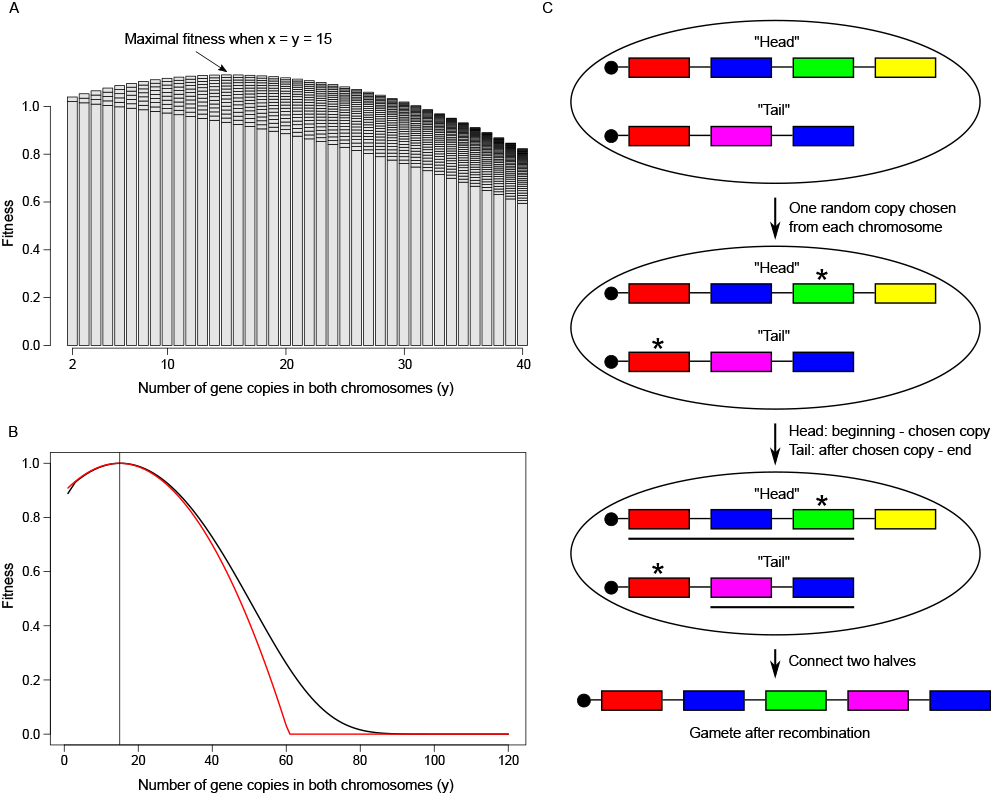
**A**. Fitness of an individual as a function of *x* (stacks) and *y* (bars). Parameters: *s*_*x*_ = 0.02, *s*_*y*_ = 0.005, *β*_*x*_ = 0.95, *β*_*y*_ = 1.05. Each bar represents one value of *y* with stacked fitness “layers” for *x* = 1 to *x* = *y*. **B**. Fitness of an individual in the *y*-only model. Parameters: *s*_*x*_ = 0.02, *s*_*y*_ = 0.005, *ε* = 0.05 (black) and its Taylor-approximated version 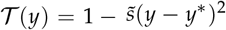, with 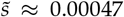 (red). The vertical line marks *y*^∗^ ≈ 14.86. **C**. Illustration of individual genotype unequal recombination. Recombination occurs in an individual with *y* = 7 = 4 + 3 gene copies and *x* = 5 < 4 + 3 different alleles (colours). The black bullet on each chromosome represents the RRM locus (see text).

The cost is only counted from the third copy, since the ground state is a single copy gene with exactly one copy on each chromosome. The selection coefficients *s*_*x*_, *s*_*y*_ > 0 and the epistasis parameters 0 < *β*_*x*_ ≤ 1 ≤ *β*_*y*_ are independent of *i*. In the following, we omit index *i* unless required for clarity. The way we define epistasis reflects the classical concepts of diminishing returns (*β*_*x*_) and synergistic epistasis (*β*_*y*_): the benefit of adding new alleles decreases with the number of already existing alleles. Think of the physiological limit preventing perfect recognition of an infinite number of possible pathogens or sensory stimuli in nature. In contrast, the cost of adding more copies increases with the number of already existing copies. This reflects the growing threat to genome integrity by inserting more and more copies.

For any fixed copy number *y*, fitness is maximized when *x* = *y*, i.e., when every copy is a different allele (which is an assumption in the *y*-only model). Whether fitness is maximized for small or for large *y* depends on the relative magnitudes of *s*_*x*_ and *s*_*y*_: assuming *x* = *y* and *s*_*x*_ ≤ *s*_*y*_, maximum fitness is achieved at the lowest possible copy number, *y* = 2. Arguably, this situation represents the standard scenario for single copy genes in nature: the cost of adding copies would outweigh its benefit. In contrast, when *s*_*x*_ > *s*_*y*_, maximum fitness may be attained at values *y* > 2. Without epistasis, and as a function of *y*, fitness is monotonically increasing, with lowest fitness at *y* = 2. With epistasis, fitness has a non-trivial maximum at *y*^∗^ (Fig. 1A). In this case, we have (see Appendix)

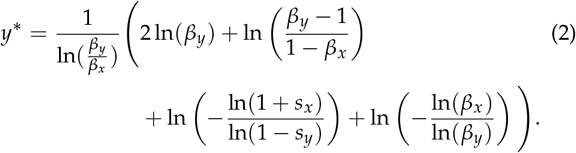

Assuming further *β*_*x*_ = 1 − *ε* and *β*_*y*_ = 1 + *ε* for small *ε* > 0, and using ln(1 + *ε*) ≈ *ε, y*^∗^ simplifies to

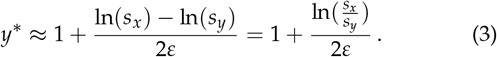

In finite populations alleles are lost by drift. Although new alleles are introduced by mutation, one generally has *x* < *y* at mutation-drift equilibrium. We employ an infinite alleles model: mutation occurs with rate *µ* per copy per individual per generation and turns a given allele into a new, previously non-existing one. The more copies an individual has, the more likely a new allele will be generated. Note that mutation does not change *y* or *y*′, but it may increase *x* and *x*′. The *y*-only model can be interpreted as the limiting scenario for large mutation rates such that any two copies are different. Therefore, mutation is explicitly required only in the simulations of the compound model, but not for the analytical results of the *y*-only model.

In both the compound and the *y*-only models recombination may be non-homologous, or *unequal*. As a consequence, copy number may change across generations. It is implemented as follows (Fig 1C): first, choose a pair of chromosomes and decide whether recombination occurs (probability *r*) or not (1 − *r*). In the first case, randomly mark a gene copy on both chromosomes. Then, the “upstream” fragment *in*cluding the marked copy of chromosome *m* (“head”), say, is fused with the “downstream” fragment *ex*cluding the marked copy of chromosome *p* (“tail”). For simplicity we assume recombination break points to lie outside of genes and exclude the possibility that genes may be disrupted by recombination. Only one recombination product is considered further. If the last copy was marked on the tail chromosome, no copy is added to the head fragment. Starting from two chromosomes with *y*^*m*^′ and *y*′^*p*^ copies, copy number in the offspring gamete can range between 1 and *y*^*m*^′ + *y*′^*p*^ − 1. More precisely, copy number in the offspring chromosome is a sum of uniform random variables with

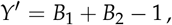

where *B*_1_ ~ U(*y*^*m*^′), *B*_2_ ~ U(*y*′^*p*^) are uniform on the integers {1, …, *y*^*m*^′}and {1, …, *y*′^*p*^}, respectively. The sum *Y*′ is trapezoidal with

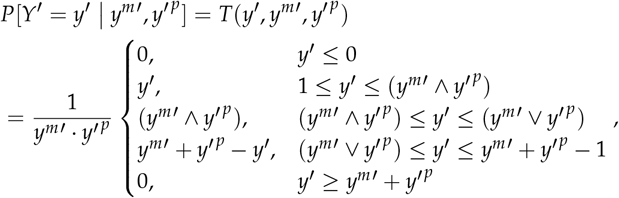

where ∧ denotes the minimum and ∨ the maximum. When no recombination occurs, only one of the two parental chromosomes is propagated.

We also consider a version with recombination rate variation: we assume that each chromosome carries a recombination rate modifier (RRM) locus which encodes a chromosome-specific recombination rate. For a pair of chromosomes *m* and *p*, a recombination event occurs with rate 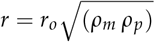 for modifier “alleles” *ρ*_*m*_, *ρ*_*p*_ > 0 which are multipliers of the base recombination rate *r*_*o*_. The modifier allele inherited to the recombination product is the geometric mean 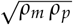. Note that selection, operating on the genotype, exerts an indirect force on the recombination rate. Symbolically, the modifier locus is represented by the black bullet in Figure 1C. It is itself not subject to recombination, but is attached to the first gene copy. We set *r*_*o*_ = 0.01 in all simulations.

### Simulations

For all simulations we used an in-house developed R programme (https://github.com/y-zheng/Recombination-gene-family) implementing a Wright-Fisher-type model with discrete generations and multinomial sampling of gametes. Simulation raw data can be downloaded from the same repository. Simulations consisted of a burn-in phase and an observation phase in which the statistics shown in Table 1 were recorded at certain time intervals. We considered four basic scenarios:

**Table 1.**
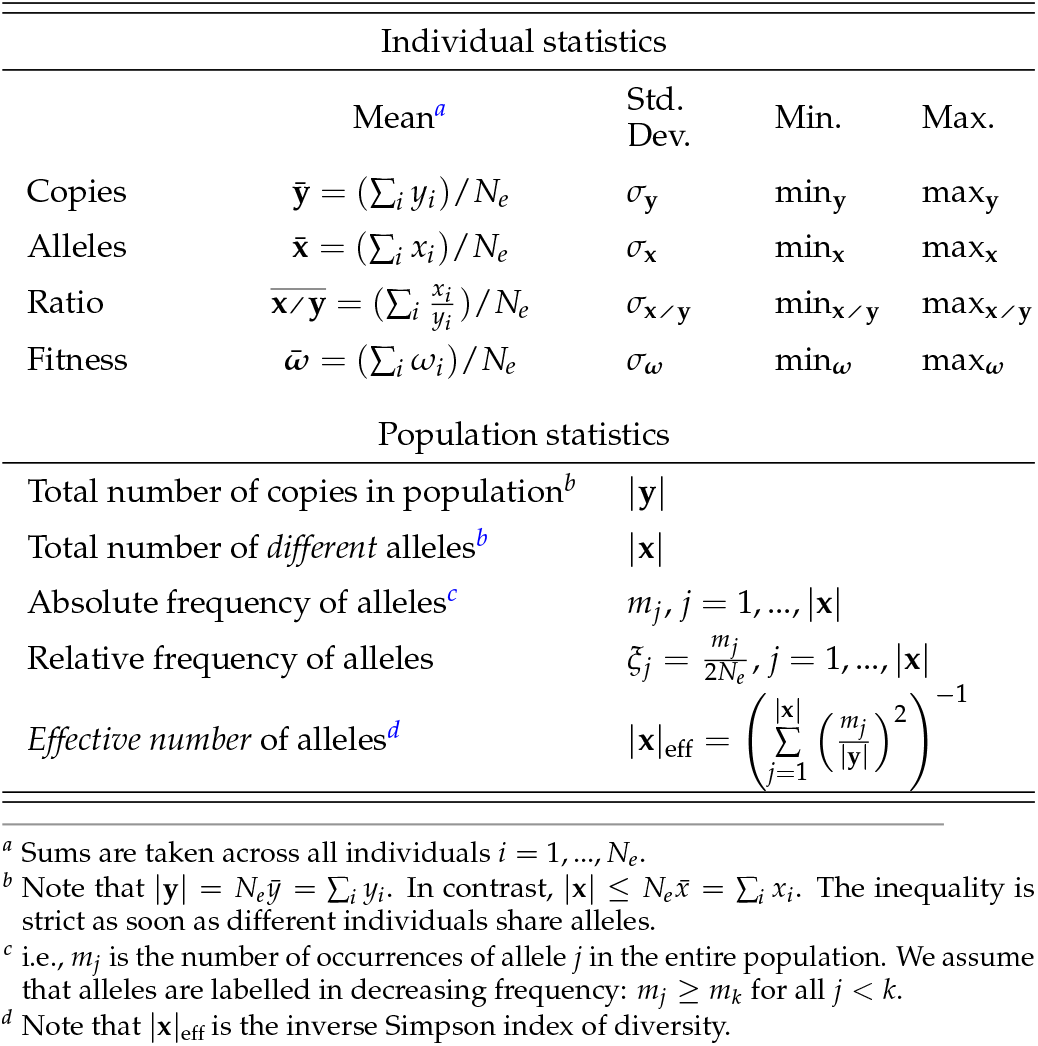
Summary statistics recorded in simulations.

a. single population with constant size *N*;
b. single population with bottleneck;
c. two sub-populations with reciprocal migration;
d. single population of constant size with RRM.

Simulations for scenario (a) were started with *y* = 10 and *x* = 1 for all *i* and run for an initial burn-in phase of 20, 000 generations. A run was re-started in case it entered during burn-in the (absorbing) state *y* = 2, i.e. when all individuals have only a single copy on each chromosome. To start simulations in scenarios (b)-(d), we used the final state which was reached at the end of scenario (a). To reduce standard error of the mean of this final sampling point, we we ran 500 replicates for scenario (a) and 200 replicates for scenarios (b)-(d). For the simulations we selected parameter ranges which we considered realistic and which turned out to be compatible with the estimates for *s*_*x*_, *s*_*y*_ and *r* and the mean copy number obtained from empirical data (see below). The parameters used in the different scenarios are listed in Table A1 in the Appendix.

### Empirical data

Based on data from the pilot phase of the 1000 Genomes project, Brahmachary *et al*. (2014) analyzed copy number variation in 193 gene families and microsatellite loci in three human populations (CEU, CHB, YRI). We chose three representative examples (`PSG3`, `MUC12` and `PRR20A`) which satisfied the following criteria:

- genes tandemly arrayed
- genes autosomal
- mean copy number between 10 and 20
- one example each with small, intermediate and large copy number variance.

`PSG3` (Pregnancy specific glycoprotein 3) is located on the long arm of the particularly gene rich chromosome 19 (Grimwood *et al*. 2004). It is a member of the carcinoembryonic antigen gene family and of the immunoglobulin superfamily and is involved in pregnancy maintenance. `MUC12` (Mucin 12) is a membrane glycoprotein of the mucin family. Mucins are involved in mucous protection, epithelial cell differentiation and intracellular signalling and have been recognized having similar evolutionary features as HLA genes (Vahdati and Wagner 2016). `PRR20A` (Proline-rich protein 20A) is a predicted gene located on the long arm of chromosome 13. It has low Uniprot annotation score with experimental evidence only at transcript level (https://www.uniprot.org/uniprot/P86496).

The available empirical data from this data set can be analyzed in the context of the *y*-only model. To estimate the underlying parameters (*s*_*x*_, *s*_*y*_, *r*) of the *y*-only model that best describe the empirical copy number distribution we implemented an *EM*-like grid search as follows: we use the data from the African (YRI) population, assuming that it is closest to recombination-selection-drift equilibrium and has highest *N*_*e*_. `PSG3`, `MUC12` and `PRR20A` have respectively mean copy numbers of 11.85, 14.94, 19.85 copies per individual in the YRI population. Individual copy numbers, as derived from the original publication (Brahmachary *et al*. 2014), can be found at https://github.com/y-zheng/Recombination-gene-family. We uniformly sample 5, 000 parameter combinations of independently chosen *s*_*x*_, *s*_*y*_ and *r* from the product of initial intervals [1e − 6, 5e − 2]^3^. For each parameter combination we calculate the Gamma-approximation of the equilibrium distribution of the *y*-only model (see Results) and use the Kolmogorov-Smirnov (KS) test to calculate the probability that the data are sampled from this distribution. We choose the top 100 (= 2%) parameter combinations to define the range of the new parameter intervals to sample from. In each iteration parameter intervals are shrinking and we terminate this process after 10 iterations to obtain a possibly small range of the final parameter combinations with highest KS *p*-value. We then chose the best parameter combinations for further analysis. Note however, that other combinations may also produce a high KS *p*-value. The range of these parameters is shown in Fig **S1**. Epistasis is kept fixed at *ε* = 0.05 during the entire search.

## Results

### y-only model

Consider first the *y*-only model. Each copy is considered a unique and distinct allele. Therefore, at any time, *x*_*i*_ = *y*_*i*_ ∀*i*, and fitness of an individual is a function only of *y*:

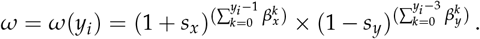

for all individuals *i*.

Let *y*′ be the number of gene copies on a single chromosome, without regard of parental status, and let *p*_*t*_ (*y*′) be the frequency of chromosomes with *y*′ copies in an infinitely large population in generation *t*.

Choosing parental chromosomes according to their fitness *ω*(*y* = *y*^*m*^′ + *y*′^*p*^), the frequency of *y*′ changes to

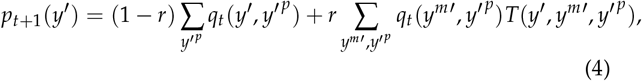

where *T* denotes the trapezoidal distribution and

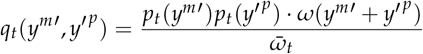

is the frequency of the pair (*y*^*m*^′, *y*′^*p*^) after selection. In the last equation 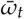 is mean population fitness at time *t*, i.e.

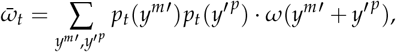

where the sum runs over all possible pairs (*y*^*m*^′, *y*′^*p*^) ∈ ℕ × ℕ. Therefore, this process can be thought of as an irreducible aperiodic Markov chain on the state space {1, 2, …}, which converges to its unique stationary distribution. Under neutrality (*ω* ≡ 1), this simplifies to

#### Proposition 1. Under (unequal) recombination and under neutrality it holds that

- *the expected value of copy number remains constant over time*, ∀_*t*_

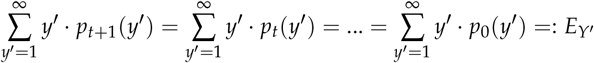
- *the stationary distribution is given by the Gamma-distribution with shape parameter α* = 2 *and expected value E*_*Y′*_, *i*.*e*.

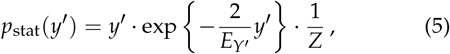

*where Z is the normalisation constant given by*

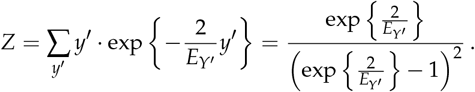

The proof is given in the Appendix.

Hence, the neutral equilibrium distribution of copy numbers on individuals is given by the convolution

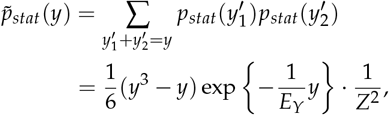

which is the Gamma distribution with shape parameter *α* = 4 and expected value *E*_*Y*_ = 2*E*_*Y′*_.

Adding selection to the process makes the analysis less straightforward. We note that the process described by equation (4) is still an irreducible Markov chain which has a stationary distribution. However, determining a closed formula of *p*_*stat*_ is not easily feasible and we resorted to the following approximation.

We choose *ω* as defined in eq. (1), assume that |**x**| = |**y**| (*y*-only model) and that *β*_*x*_ = 1 − *ε* and *β*_*y*_ = 1 + *ε* for some *ε* > 0. Thus, the fitness function simplifies to

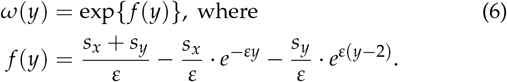

The Taylor expansion up to order 2, evaluated at *y*^∗^ and scaled with *ω*(*y*^∗^)^−1^ is

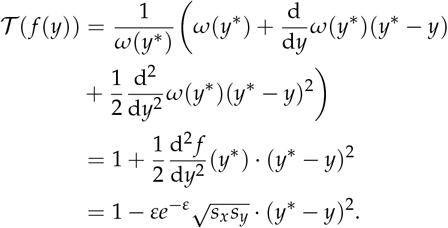

Note, that this coincides with the fitness function introduced by (Krüger and Vogel 1975)

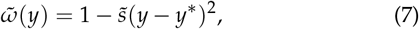

when substituting

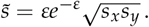

Hence, the quadratic distance of *y* from the optimal copy number *y*^∗^ determines fitness. It fits well with our definition of synergistic epistasis when *y* is not too far from *y*^∗^ (see Fig 1B) and yields a threshold 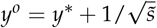 with 𝒯 (*f* (*y*)) < 0 for *y* < *y*^*0*^.

Therefore, with this quadratic approximation of the fitness function, eq. (4) becomes a finite system of equations, which can be numerically solved with standard iteration algorithms. Starting with an arbitrary initial distribution we iterate until

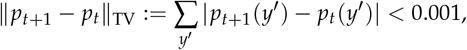

Where ‖ · ‖ _TV_ denotes the *total variation* and the sum runs from 1 to the maximal *y*′ given by *y*^*o*^. After convergence, we calculate the copy number distribution on *individuals* as convolution of the copy number distribution on chromosomes.

We find that the process converges to the same limiting distribution, independent of the initial distribution. However, different recombination rates lead to different limiting distributions, that lie between the stationary distribution under neutrality (when *r* is high), and a distribution with a sharp peak centered at *y*^∗^ (when *r* is low; with almost vanishing variance when *r* < 0.01 *s*_*x*_). When selection is strong, mean copy number of the population is shifted towards *y*^∗^. Therefore, the stationary distribution is determined by a balance of recombination and selection. Visual inspection of the limiting distribution for various parameter choices suggests that it is well approximated by a Gamma distribution also in the non-neutral case and we estimate its parameters as follows:

We numerically solved the system of equations (eq (4)) for about 50, 000 random parameter combinations. We kept *ε* = 0.05 constant and chose *r* ∈ [0, 0.01], *s*_*x*_ ∈ [0, 0.05] and *s*_*y*_ such that *s*_*x*_ /*s*_*y*_ ∈ [2.5, 18], producing an optimal copy number *y*^∗^ between 10 and 30. Then, we calculated mean and variance of the equilibrium distribution for all parameter combinations. Assuming that the expectation (*E*_*Y*_) of the limiting Gamma distribution is determined by eq (3), we set

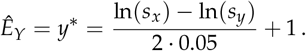

Assuming *r* > 0.01 · *s*_*x*_ and that its standard deviation scaled by the mean (*σ*/*E*_*Y*_) depends on recombination-selection balance, ln(*r*/*s*_*x*_), we obtain by linear fitting (Fig 2A):

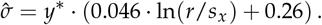

**Figure 2.**
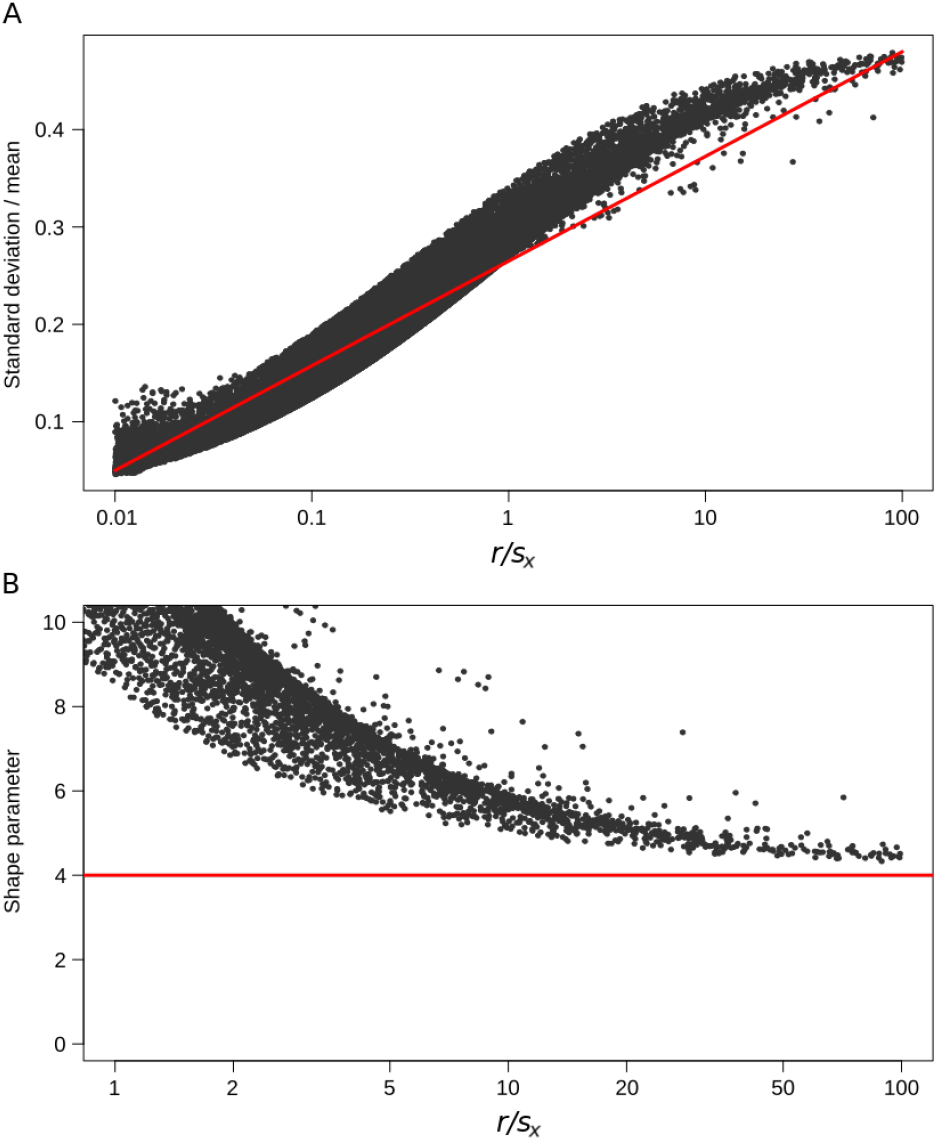
**A** Linear fit of *σ*/*E*_*Y*_ on ln(*r*/*s*_*x*_) (for details see text). Note the strong correlation of ln(*r*/*s*_*x*_) and *σ*/*E*_*Y*_, with a Pearson correlation coefficient of *ρ* = 0.97. The estimated regression line *σ*/*y*^∗^ = 0.046 · ln(*r*/*s*_*x*_) + 0.26 is shown in red. **B** Convergence of the Gamma shape parameter *α* = (*E*_*Y*_ /*σ*)^2^ towards the value *α* = 4, expected under neutrality, when *r* is increasing or *s*_*x*_ is decreasing.

Furthermore, (*E*_*Y*_ /*σ*)^2^ converges towards the shape parameter (*α* = 4) of the Gamma distribution under neutrality, when selection becomes small or recombination becomes large (Fig 2B). Therefore, for given parameters *r, s*_*x*_, *s*_*y*_ and *ε* = 0.05 we use the Gamma distribution with shape parameter 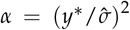 and expected value *y*^∗^ as an approximation of the equilibrium distribution of the *y*-only process with selection. Note that the distribution is uniquely determined by its shape and mean.

### Application of the *y*-only model to empirical data

To estimate selection coefficients and rates of unequal recombination for the three gene families `PSG3`, `MUC12` and `PRR20A` we used the *EM*-like grid search described above. We calculated the KS-test *p*-value for three distributions: (1) a neutral equilibrium distribution 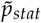 with mean value given by the arithmetic mean of the data, (2) one of the best-fitting Gamma-distributions with parameters given by the *EM*-like grid search and (3) the equilibrium distribution of the *y*-only process with the same recombination and selection coefficients as obtained from the grid search. Sufficiently small *p*-values indicate a significant difference from any of the three models, whereas a *p*-value close to one can be interpreted as a good approximation of the data. The results are given in Table 2 and Fig 3. Distributions of the 100 best parameter combinations for each gene are shown in Fig **S1**. For `PSG3` the empirical distribution of copy numbers (histogram in Fig 3, top) is well approximated by a Gamma distribution (red line) yielding a KS-test *p*-value of 0.99. The limiting distribution under the *y*-only model still fits fairly well with *p* = 0.82 (blue line). In contrast, the hypothesis of neutrality can be clearly rejected: the neutral Gamma distribution (eq. (5)) produces a *p*-value of 1.4e − 9 (black line). The parameter estimates suggest a small recombination rate of about 0.1% per generation per gamete and strong selection (*s*_*x*_ = 0.04 and *s*_*y*_ = 0.01), maintaining copy number close to its optimal value. Although the gene family `PRR20A` is much more variable than `MUC12` (Fig 3, middle and bottom) we estimate the same recombination rate of about 0.8% for both families. However, the difference in their distributions can be explained by different selection strengths. The estimates in `MUC12` are *s*_*x*_ = 0.017 and *s*_*y*_ = 0.006 – about half as strong as in `PSG3`. In contrast, the estimates in `PRR20A` are *s*_*x*_ = 0.001 and *s*_*y*_ = 0.00028, lower by roughly a factor of 40 than in `PSG3`. While neutrality can still be clearly rejected in `MUC12` (*p* = 0.0012), it cannot be rejected in `PRR20A`. Still, also for this gene family pure neutrality has a much lower explanatory power than do have models with selection (*p* = 0.217 *vs. p* = 0.98).

**Table 2.** Parameter estimates for empirical data obtained by *EM* grid search.

| Gene family | Estimated parameters | p-value of KS-test |  |  |
| --- | --- | --- | --- | --- |
|  |  | Neutral | Gamma | y-only |
| PSG3 | $r = 0.001$ | $1.4e-9$ | 0.99 | 0.82 |
| | $s_x = 0.04$ | | | |
| | $s_y = 0.01$ | | | |
| MUC12 | $r = 0.008$ | 0.0012 | 0.99 | 0.98 |
| | $s_x = 0.017$ | | | |
| | $s_y = 0.006$ | | | |
| PRR20A | $r = 0.008$ | 0.217 | 0.98 | 0.98 |
| | $s_x = 0.001$ | | | |
| | $s_y = 0.00028$ | | | |

**Figure 3.**
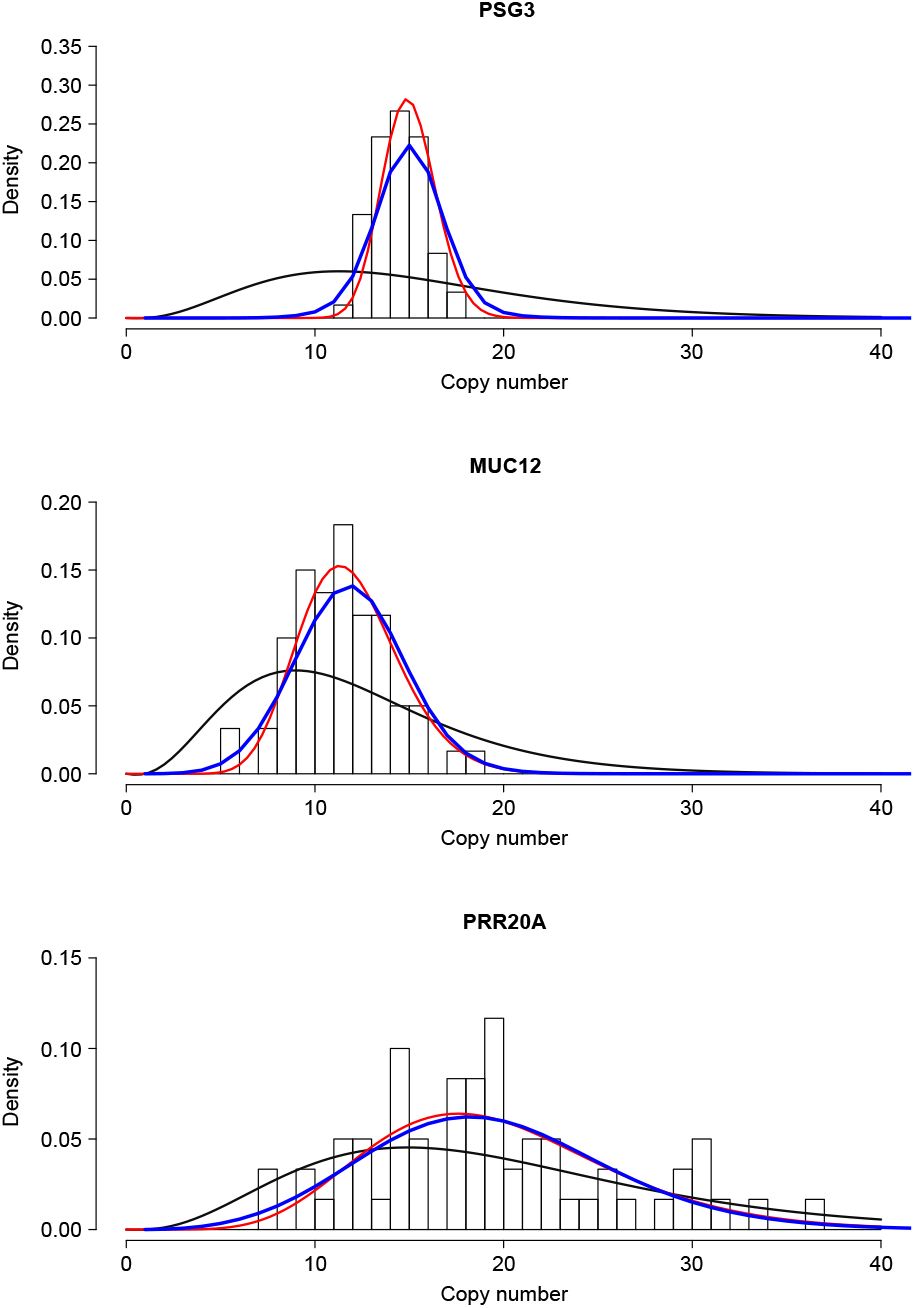
The distribution of copy number in the y-only model under neutrality (black), with Gamma-approximation (red) and equilibrium distribution of the y-only model with selection (blue).

### Simulation results of the compound model

In **scenario (a)** we analyzed the effect of different population sizes, selection strengths **(a1)** and recombination rates **(a2)** on the statistics of Table 1 at equilibrium. In **scenario (a1)** we used *s*_*x*_ = 0.01, 0.02, 0.04 (weak, medium and strong selection), with *s*_*x*_ /*s*_*y*_ = 4 and *ε* = 0.05. These parameters were chosen such that the optimal genotype for an individual is *x* = *y* = 15 in all three selection regimes. Population size varied from *N*_*e*_ = 500, 1000, 2000 to 4000 and recombination rate was kept constant at *r* = 0.01. Results are shown in Fig 4.

**Figure 4.**
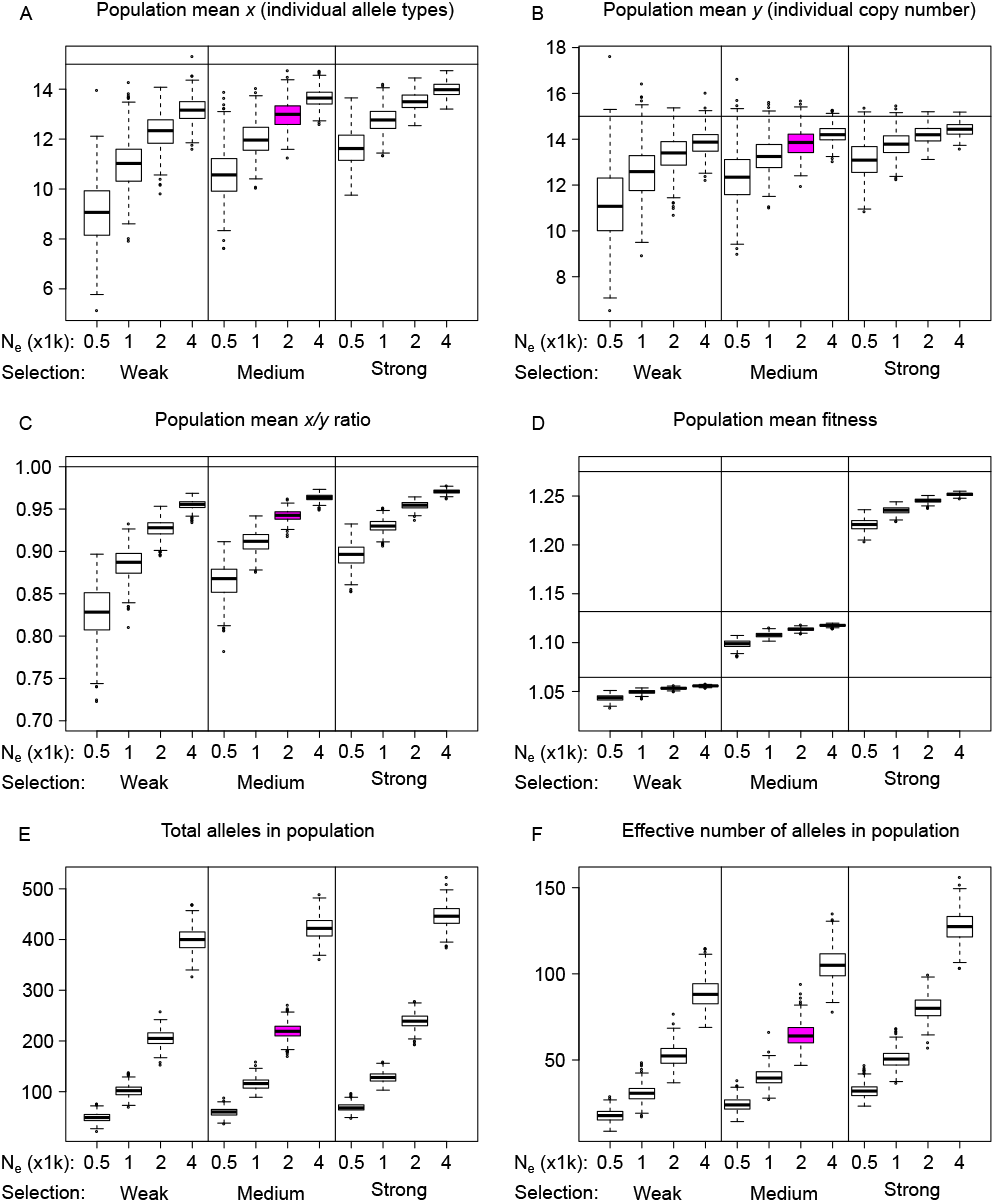
**Scenario (a1)** – constant population size. Population statistics at equilibrium: population mean 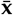 (A); population mean 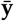 (B); 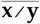 ratio (C); population mean fitness (D); total number (E) and effective number of alleles |**x**| _eff_ (F). Varying parameters: population size *N*_*e*_ and selection coefficient *s*_*x*_. Mutation (*µ* = 0.0005) and recombination rate (*r* = 0.01) are kept fixed. Boxplots based on 500 independent replicates. Box coloured in purple indicates a parameter combination (*N*_*e*_ = 2000, *r* = 0.01, *s*_*x*_ = 0.02, *s*_*y*_ = 0.005) shared by scenarios (a), (b), (c) and (d). Horizontal lines in A-C indicate the optimal copy number in the *y*-only model. Horizontal lines in D indicate optimal fitness.

Both larger population sizes and stronger selection lead to an increase in population means 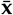 and 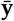 (Fig 4A and B). Note, that the demographic effect (decrease of drift by increase of population size) on these quantities is much stronger than the effect by increasing selection. Both 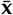 and 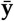 are always below the optimal value of 15. However, doubling *N*_*e*_ has a stronger effect than doubling selection strength in bringing the population closer to the optimal value. Essentially the same pattern is observed for the ratio 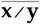 (Fig 4C). For example, *N*_*e*_ = 1000, 2000, 4000 with low selection leads to a higher ratio 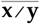 than *N*_*e*_ = 500, 1000, 2000 with intermediate selection. The total (Fig 4E) and the effective (Fig 4F) number of alleles scale roughly linearly with *N*_*e*_. Again, both quantities depend more strongly on population size than on selection strength. This effect is more pronounced in the total number of alleles than in |**x**|_eff_, which is explained by drift: alleles at low frequency, in particular newly generated alleles (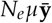 per generation), are prone to loss when drift is strong. They count for the total number, but contribute little to |**x**|_eff_. In contrast, mean fitness is strongly affected by the strength of selection: *s*_*x*_ has a much larger effect than *N*_*e*_ (Fig 4D). Finally, the frequencies of the most common alleles (Fig **S2**) are negatively correlated both with *N*_*e*_ and *s*_*x*_. Summarizing, allelic diversity at population scale appears to be driven mainly by *N*_*e*_.

In **scenario (a2)** we kept selection at intermediate level (*s*_*x*_ = 0.02, *s*_*y*_ = 0.005) and varied the rate of (unequal) recombination from *r* = 0.002 to 0.05. Results are shown in Fig 5. Increasing recombination decreases 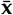 and 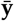, as well as the ratio 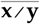. Therefore, it also decreases mean fitness 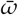. Recombination acts here in a similar way as drift: doubling the recombination rate has the same effect on fitness as halving the population size. This observation can be interpreted as a recombination load: frequent recombination can generate chromosomes whose copy number is far away from the optimum. Deviation from the optimal copy number has an asymmetric effect because of epistasis: a surplus of copies is more harmful than a deficit (Fig 1B), explaining the somewhat counter-intuitive effect that increasing the recombination rate decreases both total and effective number of alleles in the population.

**Figure 5.**
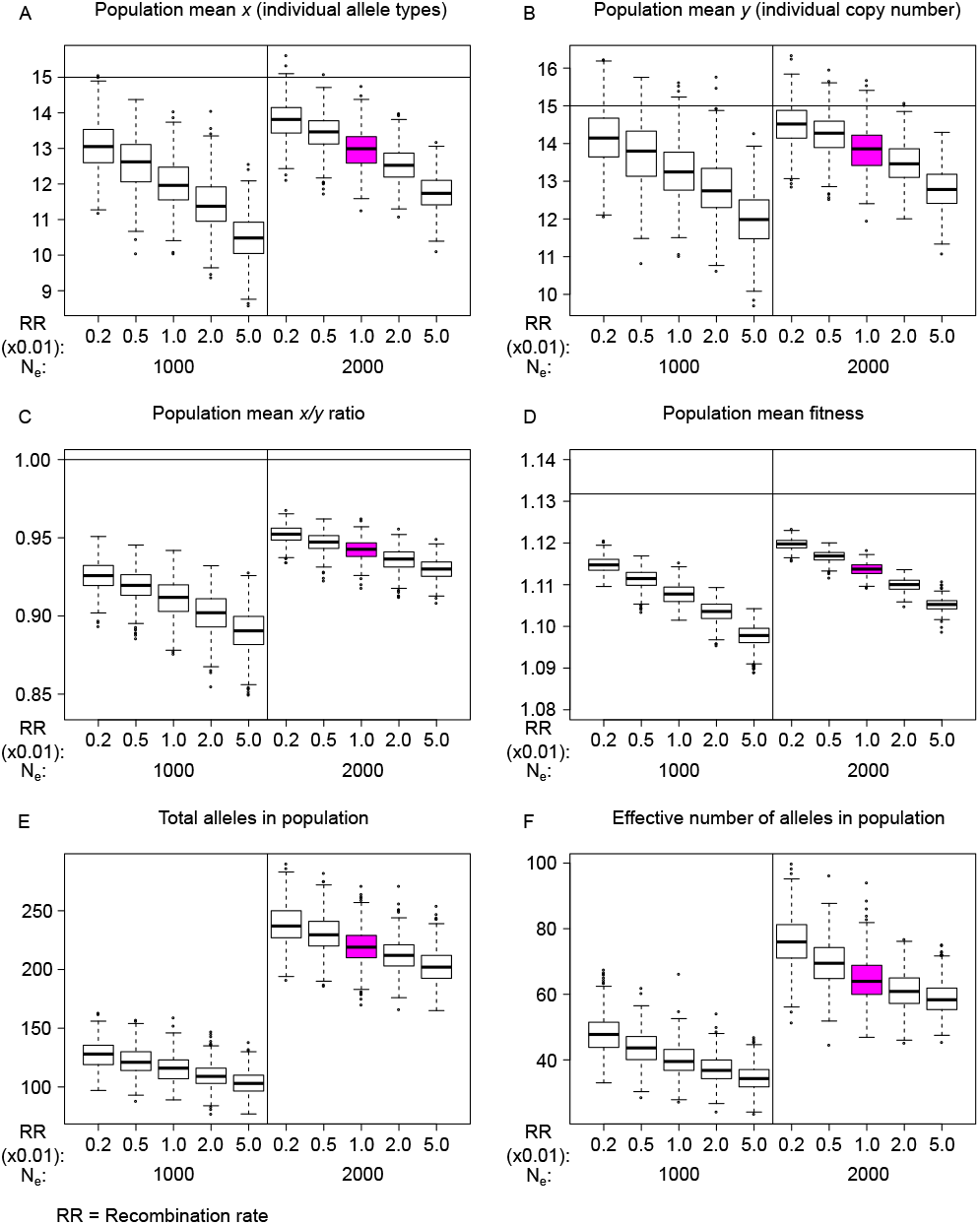
**Scenario (a2)** – constant population size. Population statistics at equilibrium: population mean 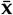 (A); population mean 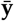 (B); 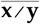 ratio (C); population mean fitness (D); total (E) and effective number |**x**|_eff_ (F) of alleles. Varying parameters: population size *N*_*e*_ = 1000, 2000 and recombination rate (*r* = 0.01 times the factor indicated on the abscissa). Mutation rate (*µ* = 0.0005) and selection strength ((*s*_*x*_, *s*_*y*_) = (0.02, 0.005)) are kept fixed. Boxplots based on 500 independent replicates. Box coloured in purple indicates the parameter combination (see Fig 4) shared by scenarios (a), (b), (c) and (d). Horizontal lines as explained in Fig 4.

In **scenario (b)** we explored the impact of a single instantaneous and short bottleneck. Starting with an equilibrated panmictic population of constant size *N* = 2000, population size was reduced to 1% (= 20) for 5, 10, or 20 generations, then restored to its original value *N* and the generation counter reset to *t* = 0. After that, simulations are carried on for another 10, 000 generations during which the recovery process of the six summary statistics mentioned above is recorded. Results for different selection strengths are summarized in Fig 6. A longer period of population size reduction results in populations with lower 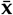 and lower 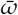. In contrast, length of the reduction period hardly affects 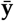. Recovery time correlates positively with the length of the reduction period.

We observed that 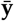 and, to a lesser extent, 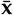 experience a decrease *after* the restoration of population size, and before it returns to its constant equilibrium value. Furthermore, the total number of alleles recovers much faster than the effective number. The reason is that new alleles are quickly created by mutation, but – while rare – they continue to bias the effective number of alleles, before equilibrium frequencies are restored. By segmental regression we found that mean fitness recovers faster than |**x**|_eff_ (Fig **S3** A and B). Furthermore, populations under stronger selection recover faster. The variation of these statistics among replicates is shown in Fig **S4**. Except for total and effective number of alleles, all other statistics show little among-replicate-variation after about 500 to 1000 generations after the bottleneck. Variation of the total number of alleles reaches a plateau and then gradually decreases, while among-replicate-variation of |**x**|_eff_ is generally small.

**Figure 6.**
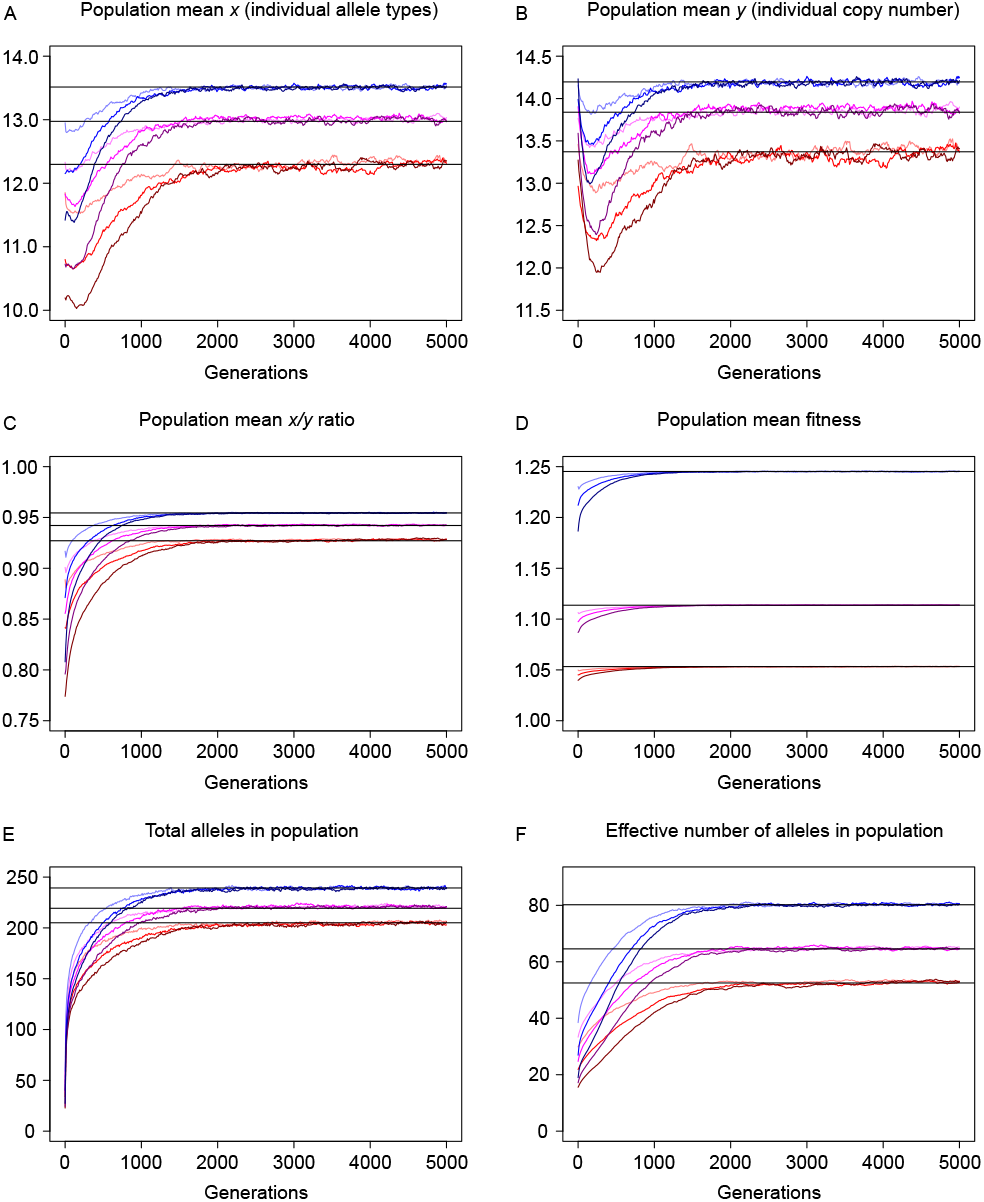
**Scenario (b)** – recovery after a bottleneck. Equilibrium populations with *N* = 2000 are reduced to *N*_red_ = 20 for a period of 5, 10 or 20 generations and then restored. During recovery six statistics are traced. A: population mean 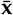; B: population mean 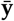; C: ratio 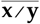; D: mean fitness 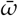; E: total number of alleles; F: |**x**|_eff_. Red, purple and blue colours indicate weak, intermediate and strong selection. Light to dark shading indicates short to long bottleneck durations. Each curve is an average across 200 replicates. Horizontal gray lines are equilibria under constant population size.

In **scenario (c)**, we studied the effect of population subdivision and migration. We simulated reciprocal migration with two sub-populations of equal size, small (*N* = 500) and intermediate (*N* = 1000), starting from pairs of independent equilibrated replicates from scenario (a). Then, time was reset to *t* = 0 and migration was turned on with rates *Nm* = 0.1, 1 or 10 individuals per generation per direction. Summary statistics 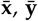, mean fitness 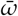, total number of alleles and |**x**|_eff_ in the combined super-population were recorded over time. After about 1500 to 2000 generations, these statistics approached a migration-drift-selection equilibrium, which is between the means for the panmictic populations of size *N*_*e*_ = 1000 and *N*_*e*_ = 2000. While the scenario with high migration (*Nm* = 10) is almost indistinguishable from the panmictic population with respect to 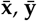 and 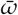 (Fig 7A-D), there is still a clear deficit in the total and effective number of alleles compared to the panmictic population, even when the migration rate is high (Fig 7E,F). Note also in this case the initial overshooting of the panmictic equilibrium in the statistics 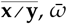 and |**x**|_eff_ at about 100-200 generations, which is reminiscent of transient “hybrid vigour”. Variation of these statistics among population replicates does not change appreciably with time (Fig **S5**). Similar results are observed for small populations *N*_*e*_ = 500 (Fig **S6** and **S7**).

**Figure 7.**
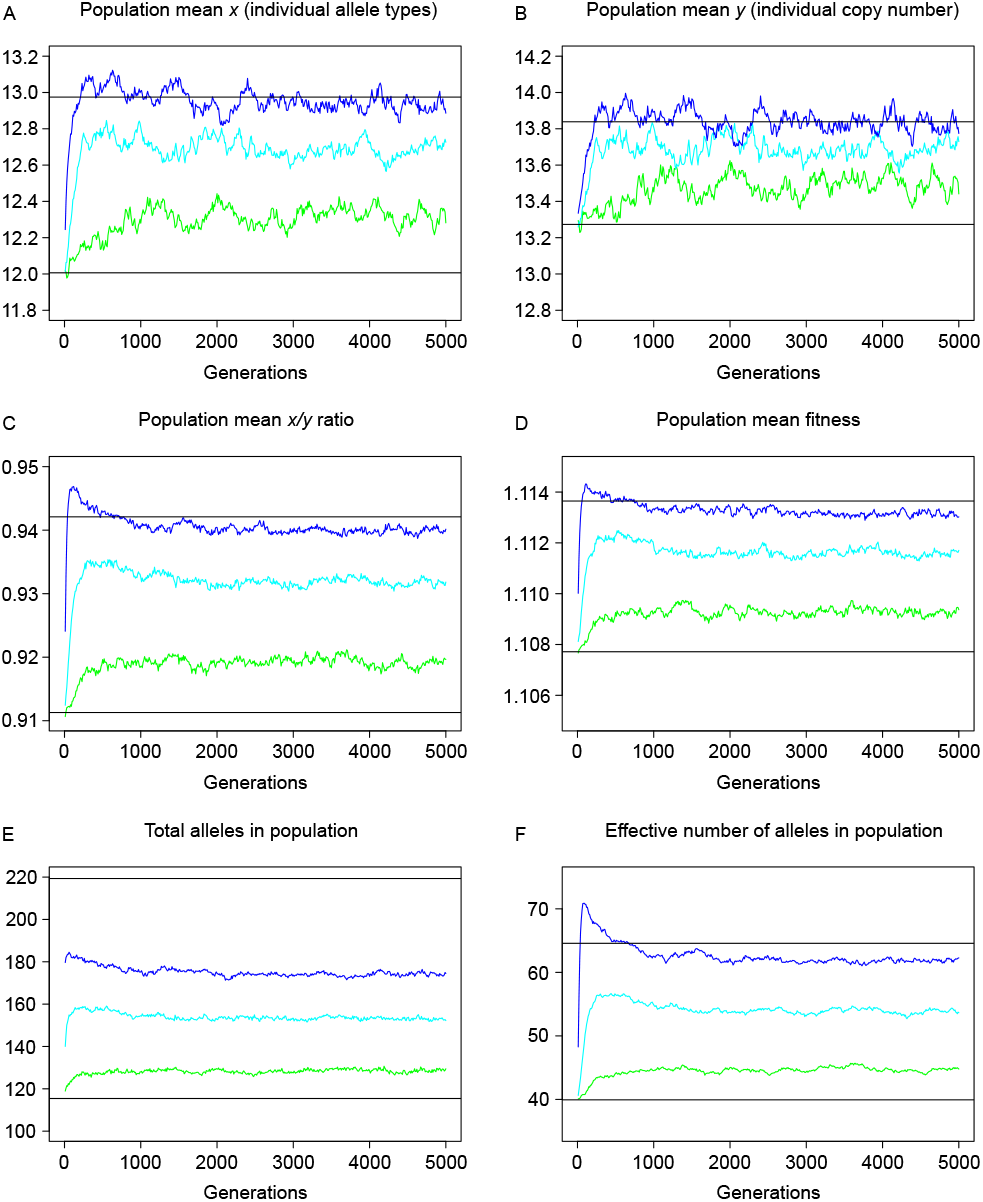
**Scenario (c)** – migration. Two separated and equilibrated sub-populations of size *N* = 1000 start to exchange migrants at time *t* = 0. Medium strength of selection (*s*_*x*_ = 0.02, *s*_*y*_ = 0.005). Migration rate: 2*Nm* = 0.1 (green), 1 (cyan) or 10 (blue) migrants per generation in each direction. (A) population mean 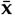; (B) population mean 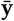; (C) ratio 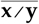; (D) population mean fitness 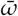; (E) total and (F) effective number of alleles in the combined super-population. Shown are mean values across 100 replicates. Black lines indicate mean values (across 500 replicates) in panmictic populations of size *N*_*e*_ = 1000 (lower line) and *N*_*e*_ = 2000 (upper line).

In scenario (a2) we observed that lower recombination rates lead to an equilibrium of 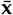 and 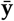 which are closer to the optimum. A natural question to ask is whether the recombination rate itself maybe subject to selection. Therefore, in **scenario (d)** a recombination rate modifier was added to the simple model. Given an equilibrated population which was reached with *r* = 0.01 as described in scenario (a), recombination rate modification was switched on, and time reset to *t* = 0. Recombination rate was coded by an RRM allele, which can increase or decrease the current recombination rate by a factor *e*^±0.05^ when mutated. Modification happens per chromosome per generation each with probability *p* = 0.0002 for increase or for decrease. The RRM locus is thought to reside on the tip of a chromosome without itself being affected by recombination (Fig 1). Simulations were carried on for 50, 000 generations and runs for each parameter setting of (*s*_*x*_, *s*_*y*_) were replicated 200 times. The results show that the mean recombination rate (average across all RRM alleles in the population) is continuously decreasing (Fig 8). It decreases more and faster when selection (*s*_*x*_ and *s*_*y*_) is strong. When simulations terminated, the recombination rate was reduced – on average – to 56%, 41% and 31% of its original value (*r* = 0.01) and it showed a strongly negative correlation with population mean fitness (Pearsons’s *r* =− 0.75, −0.83, −0.78) for weak, intermediate and strong selection, respectively.

**Figure 8.**
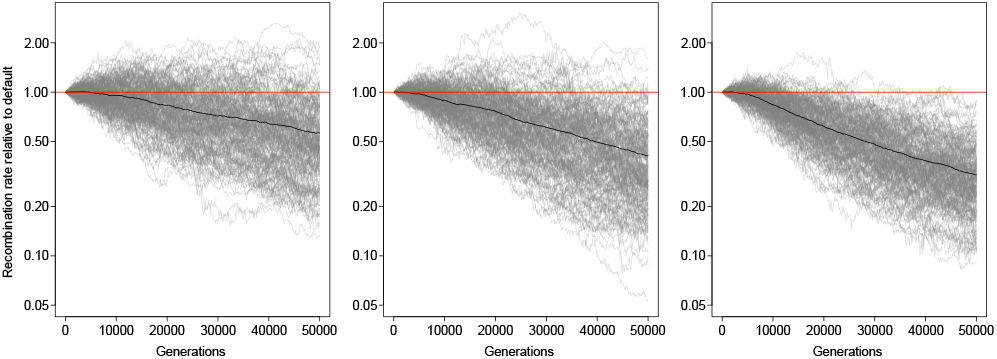
**Scenario (d)** – RRM: recombination rate modification. Populations, which have reached equilibrium without RRM, are carried on for 50, 000 generations during which the recombination rate, encoded at a modifier locus, may change under the influence of selection. For all iterations: *N*_*e*_ = 2000, *r* = 0.01. Left: weak (*s*_*x*_, *s*_*y*_) = (0.01, 0.0025); middle: intermediate (0.02, 0.005); right: strong selection (0.04, 0.01). Shown are trajectories of the recombination rate (in percent of its original value *r* = 0.01) for 200 replicates each. The mean across all 200 replicates is shown as a black line.

## Discussion

We considered here a model in which two mechanisms, unequal recombination and mutation, may generate chromosomal diversity. While mutation leads to genetic diversity *sensu strictu*, by unequal recombination a chromosome may receive additional, or lose existing gene copies. Therefore, it is similar, but not identical, to segmental duplication or loss: copies gained by unequal recombination have their origin in a pairing haplotype, hence may be genetically diverse upon arrival, while those gained by duplication have their origin in the same haplotype, hence are genetically identical upon arrival. However, this distinction is negligible, since a single mutation event already suffices to make two identical copies distinct from each other when working in the context of the infinite alleles model. Another feature of our model are the two overlaid components of the fitness function: it decreases with copy number, but increases with allele number, entailing a subtle and very interesting interaction of recombination and selection.

To gain some analytical insight into copy number dynamics under recombination, we first considered the neutral case in an infinitely large population. We find copy number of individuals to be Gamma distributed with an equilibrium mean which is identical to the initial mean at time *t* = 0 and remains constant over time. The limiting shape parameter is *α* = 4, which is identical for all initial configurations. These two properties together uniquely determine the limiting distribution, which is independent of the shape of the initial distribution and of the recombination and mutation rates.

Adding selection changes the game. The limiting distribution becomes dependent on both the recombination rate and the strength of selection, but independent from the initial configuration. Still, it is well approximated by a Gamma distribution. The distribution that results from low selection strength or high recombination converges to the neutral equilibrium. A characteristic property is the negative correlation between limiting mean copy number and recombination rate: lower recombination rate leads to a higher mean copy number at equilibrium (Fig 8).

Note, that compound fitness, in which allele diversity is credited, contains a component of balancing selection: an individual which is heterozygous at any given locus has a higher fitness than one which is homozygous at the same locus. An important difference between the model considered here and one-locus models of balancing selection is the existence of gene copy number variation and unequal recombination. Note, that allelic diversity in the population can be stably maintained even in the case of allele fixation at single loci. The possibility to maintain allelic diversity through gene duplication, or unequal recombination, has been suggested by (Haldane 1937). It is somewhat surprising that Haldane’s idea has received only little attention in classical population genetics theory nor in experimental work. To our knowledge, tests confirming Haldane’s hypothesis were conducted only a few years ago (Milesi *et al*. 2017).

We have shown that a high recombination rate has a negative effect on allelic diversity and resultant mean fitness. The reason is twofold: (1) a higher rate of unequal recombination produces individuals with much higher or lower copy number than the optimum, which have reduced fitness; (2) low recombination increases the likelihood for highly unfit homozygotes to appear, thus improving the efficiency of selection.

Populations which experienced strong bottlenecks are at risk of inbreeding depression, and loci under balancing selection are particularly affected (Frankham *et al*. 2014). Random loss of alleles increases homozygosity and consequently reduces fitness. This can affect and delay the recovery of genetic diversity even after population size has recovered (Miller and Lambert 2004). In this study, we explored the effect of some parameters on the speed and process of bottleneck recovery at loci under diversifying selection. Both selection strength and bottleneck length influence the process. Relatively longer bottlenecks produce a temporary reduction in 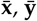 and mean fitness. The most likely reason is that high homozygosity results in selection towards haplotypes with fewer copies. Selection is more powerful after, than during, the bottleneck,when population size has recovered, but copy number recovery may lag behind. However, this somewhat paradoxical effect of fitness reduction at the initial phase of bottleneck recovery is only a short term effect, and – at least in part – due to the instantaneous, rather than gradual, restoration of population size in our model. Compared to fitness, |**x**|_eff_ is recovering even more slowly: for fitness to recover it suffices that new alleles appear and survive. But |**x**|_eff_ has recovered only when allele frequencies have reached their equilibrium values. Therefore, |**x**|_eff_ is a more sensitive statistic to test for deviation from equilibrium.

Simulations of scenario (c) show that fitness under population subdivision with moderate migration reaches an equilibrium which is intermediate between those under panmixis on the one hand and complete isolation on the other. While a short boost of hybrid vigour exists, we do not see a positive effect from limiting migration compared to panmixis. An earlier simulation study (Schierup *et al*. 2000) showed that the allelic diversity is largely insensitive to migration rates, but low-migration scenarios result in alleles with more divergent sequences. Additionally, balancing selection in the form of heterosis could increase the effective migration rate because migrant haplotypes are more likely to be successful in this case than under neutrality (Ingvarsson and Whitlock 2000). Diversifying selection on MHC alleles has been shown to increase divergence between subpopulations, while diversity within subpopulations is still mostly governed by drift (Herdegen *et al*. 2014). MHC alleles and genes are also known to be shared among species through introgression, leading to restoration of diversity previously lost by drift (Dudek *et al*. 2019). In addition to generic balancing selection also local adaptation, i.e. the fixation of alleles which are adapted to specific subpopulations, may increase allelic diversity between populations (Ekblom *et al*. 2007). However, this effect is not considered in the model presented here, where selection operates only on the number of distinct alleles.

When the recombination rate is allowed to change over time we observe a trend towards lower rates. It is driven by selection and happens on a realistic population genetic timescale of some thousand generations. However, there is little empirical knowledge about (unequal) recombination rates in multi gene families. For example, in the human MHC locus the recombination rate is only about a third of the average genomic background rate (de Bakker *et al*. 2006; Traherne 2008). On the other hand, studies on bovids (Schaschl *et al*. 2006) and horse (Beeson *et al*. 2019) show the opposite: high recombination in the MHC and olfactory receptor loci. In contrast again, the values reported for chicken seem to depend on mapping methodology (Fulton *et al*. 2016). Results from sheep (Petit *et al*. 2017) suggest a high “historical” (estimated from population data), but a low “meiotic” (from pedigree data) recombination rate, which suggests a recent change in time. From humans again, it is well known that recombination hotspots have a very fast turn-over time and are distinct in different subpopulations (Lam *et al*. 2013). Also, recombination rates may substantially differ in females and males – one example is the long arm of human chromosome 19 (Grimwood *et al*. 2004). Additionally, the presence of gene conversion makes the estimation of (reciprocal) recombination rates difficult (Martinsohn *et al*. 1999; Hosomichi *et al*. 2008). Anyway, current experimental results do not reveal a consistent picture as to whether there is a benefit, or trend, to suppress recombination in large multi gene families.

### Caveats and future direction

While our model has incorporated multiple genetic processes, it is likely still far away from the details of how multi gene families evolve in real-life populations. One issue, not considered here, is gene conversion where an allele, or a fragment thereof, overwrites another one in a pairing chromosome. For example, gene conversion is known to play an important role in maintaining MHC diversity (Martinsohn *et al*. 1999; Högstrand and Böhme 1999; Wiehe *et al*. 2000; Bahr and Wilson 2012).

Also, our selection model assumes time-independent fitness and each allele provides the same selective benefit. This corresponds to an ideal situation where external factors are ubiquitous and stable. In practice, however, the selective benefits of certain alleles do change together with a changing environment. Evolving pathogens, for instance, lead to an arbitrarily complex co-evolution dynamics (Ejsmond and Radwan 2009; Tellier *et al*. 2014). Furthermore, population structure may interact with diversifying selection in a complex or even counter-intuitive way. In humans it is known that different populations harbour different MHC alleles, likely driven by pathogen diversity (Manczinger *et al*. 2019). A hypothesis is that multiple subpopulations act as reservoirs of alleles and backups for each other, allowing for quick response against new pathogens (Lenz *et al*. 2009; Linnenbrink *et al*. 2018). Interaction between population structure and local adaptation needs to take into account subpopulation sizes and migration networks. For instance it was shown that subpopulation sizes can affect local allelic diversity (Mason *et al*. 2009).

Finally, and perhaps most importantly, gene function decides on fitness. On population genetic time scales pseudogenization plays an important role for the evolution of multi gene families (Hess 2000; Menashe *et al*. 2006). Although eventually removed by selection, pseudogenes can persist in real-life populations with high frequency. Conditions under which pseudogenes appear and persist can be identified in accordingly modified models. Structural and functional aspects being included together with gene conversion, temporally or locally varying selection strengths into theoretical models will help to address open questions, but remains to be considered in future work.

## Appendix

*Proof of (2)*. Using the closed form formula of the geometric series and the fact that *x* = *y*, we can write the fitness function 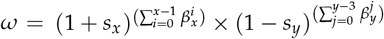 as a function of *y* that equals

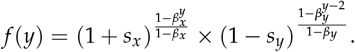

Defining

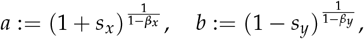

we find that

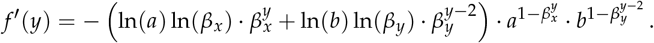

Setting *f*′ (*y*^∗^) = 0 leads us to

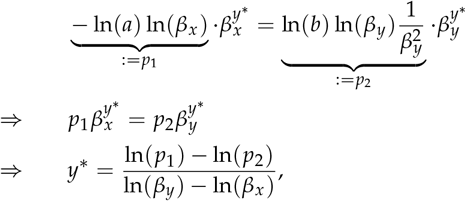

and inserting the expressions for *p*_1_, *p*_2_, *a, b* gives the result.

□

*Proof of Proposition 1*. We note that the parental status of the chromosomes do not matter in the following calculations. Therefore, we use the notation 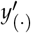 instead of *y*^*m*^′ and *y*′^*p*^. Since the *T* describes the distribution of the sum of two uniform random variables, we observe that the expected value is given by

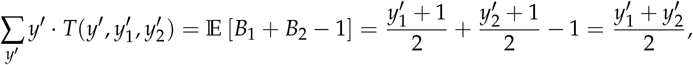

and therefore conclude that

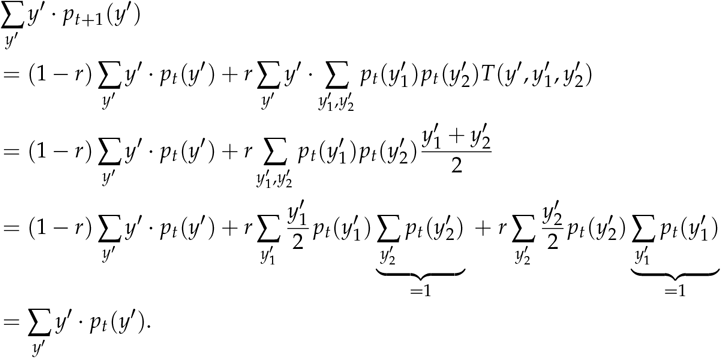

We define *a* = 2/*E*_*Y′*_ and note that the stationary distribution is independent from the recombination rate *r* > 0, i.e.

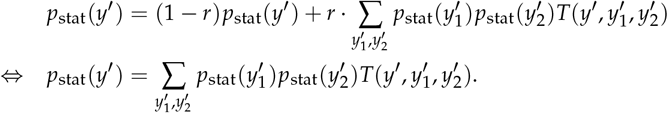

Therefore, we find that

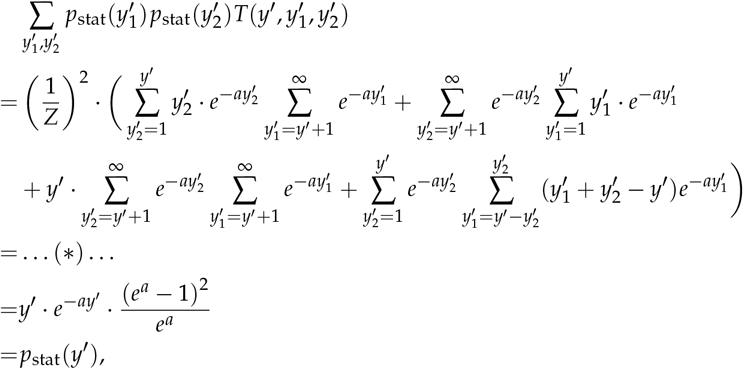

where the detailed calculations of (∗) are shown below. □

*Proof of (*)*. Using the substitution 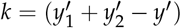 we find that

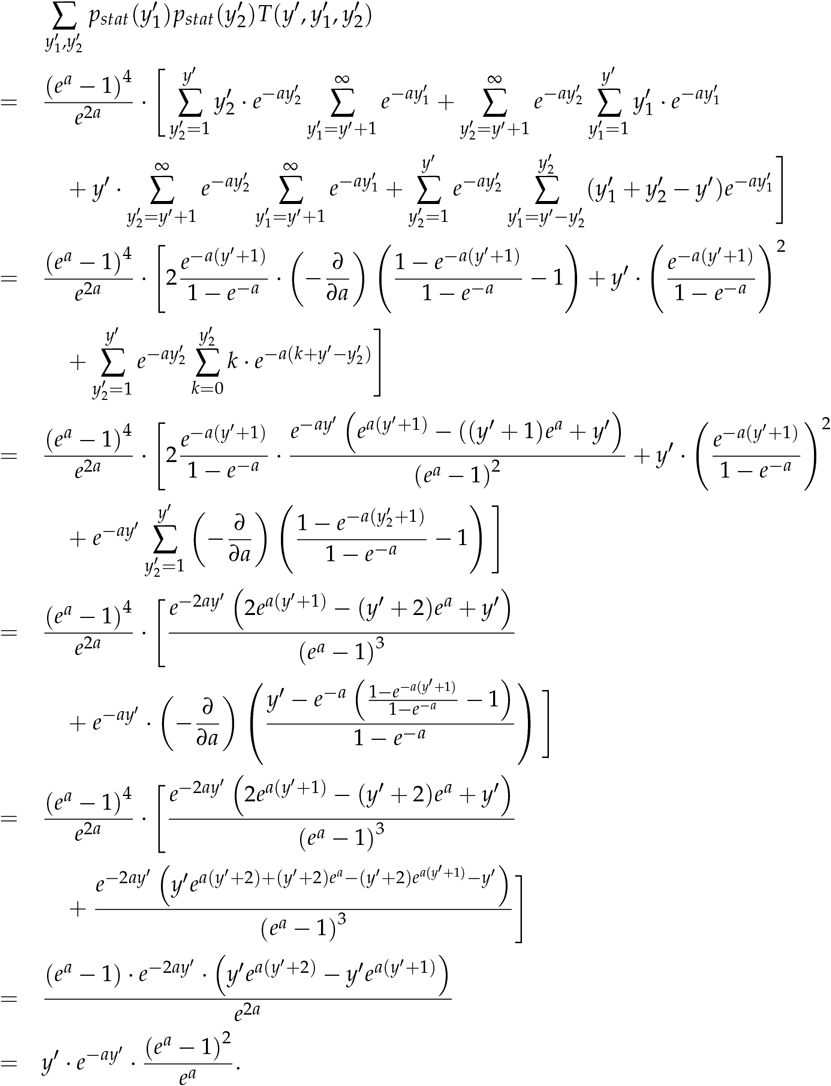

□

**Table A1.**
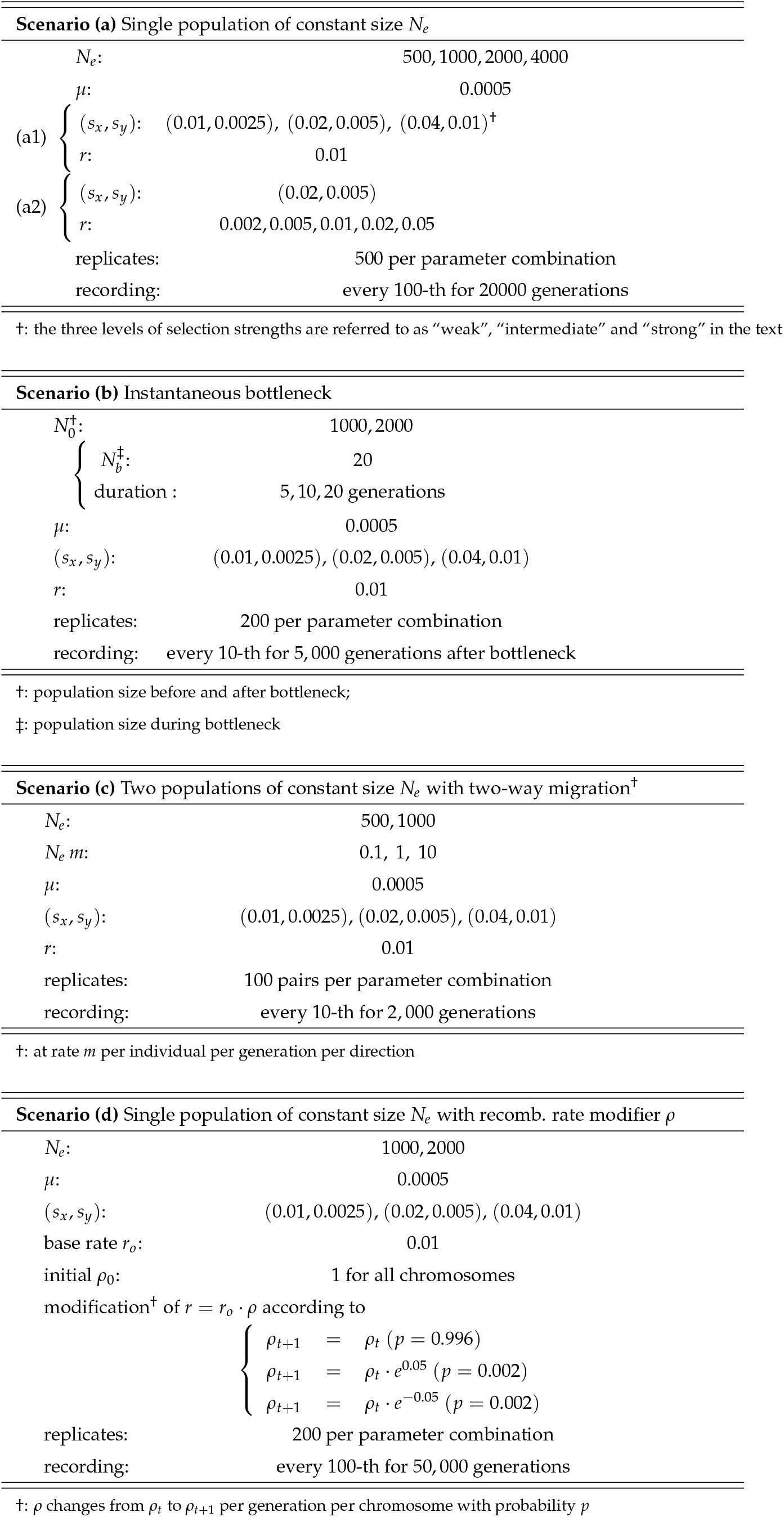
Parameters used in simulations of the compound model.

**Figure S1.**
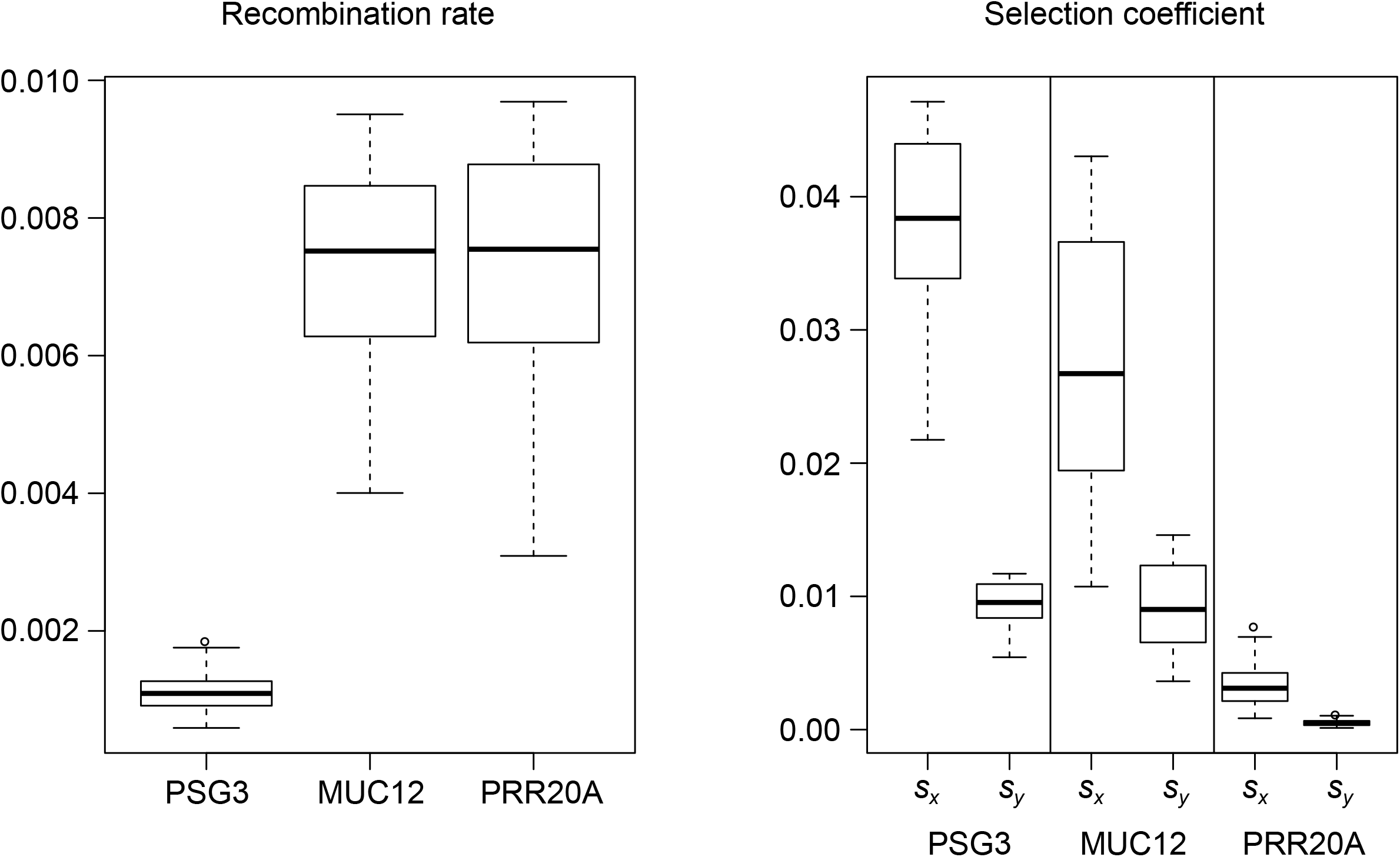
Distributions of the best 100 parameter combinations (*r, s*_*x*_ and *s*_*y*_) estimated from empirical distribution of gene copy numbers in YRI population (data from Brahmachary et al. 2014), based on the y-only model with selection. Distributions are shown separately for *r, s*_*x*_ and *s*_*y*_, although they are estimated jointly.

**Figure S2.**
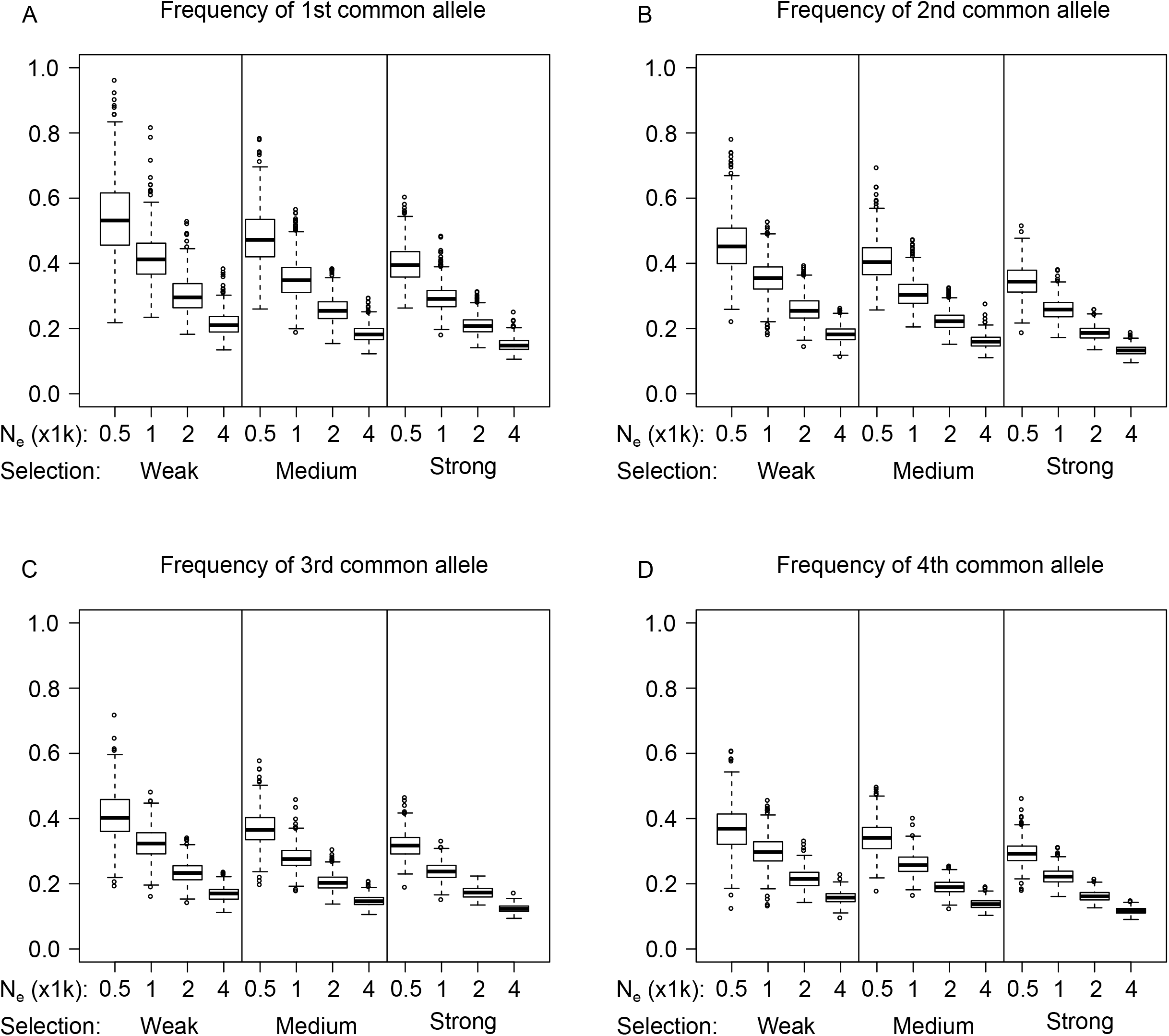
Scenario (a1) -- Frequencies *ξ*_*i*_ (*i=1*,…,*4*) of the four most common alleles (A. 1st, B. 2nd, C. 3rd, D. 4th most common) in the equilibriated population in scenario (a1). Frequency is defined as proportion of all *2N*_*e*_ chromosomes that contain the allele.

**Figure S3.**
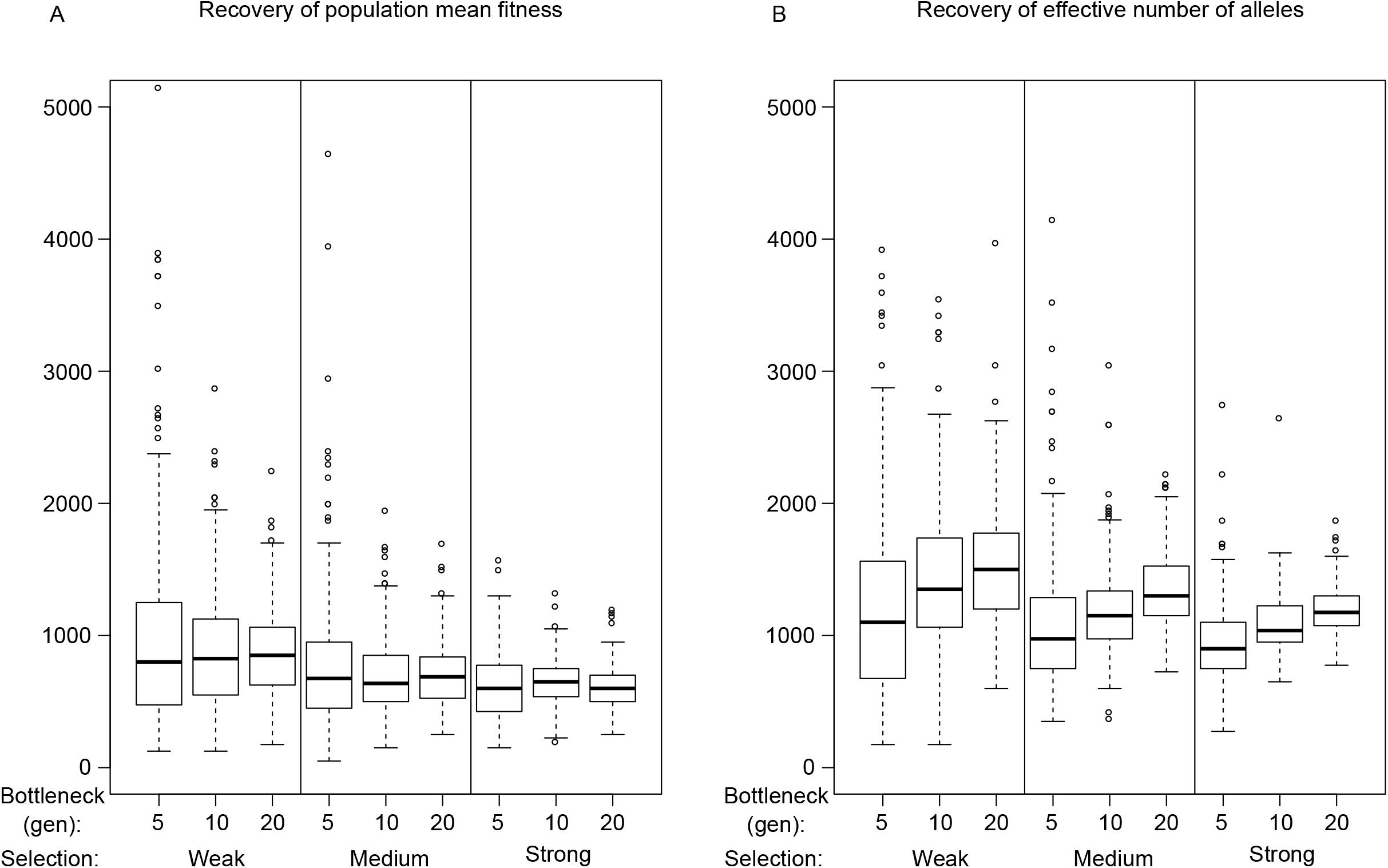
Time for A. population mean fitness, B. effective number of alleles, to return to the pre-bottleneck level, as estimated using segmental regression. The statistic is fitted to a regression line that increases linearly and becomes constant at a time point, which is used as the time for recovery.

**Figure S4.**
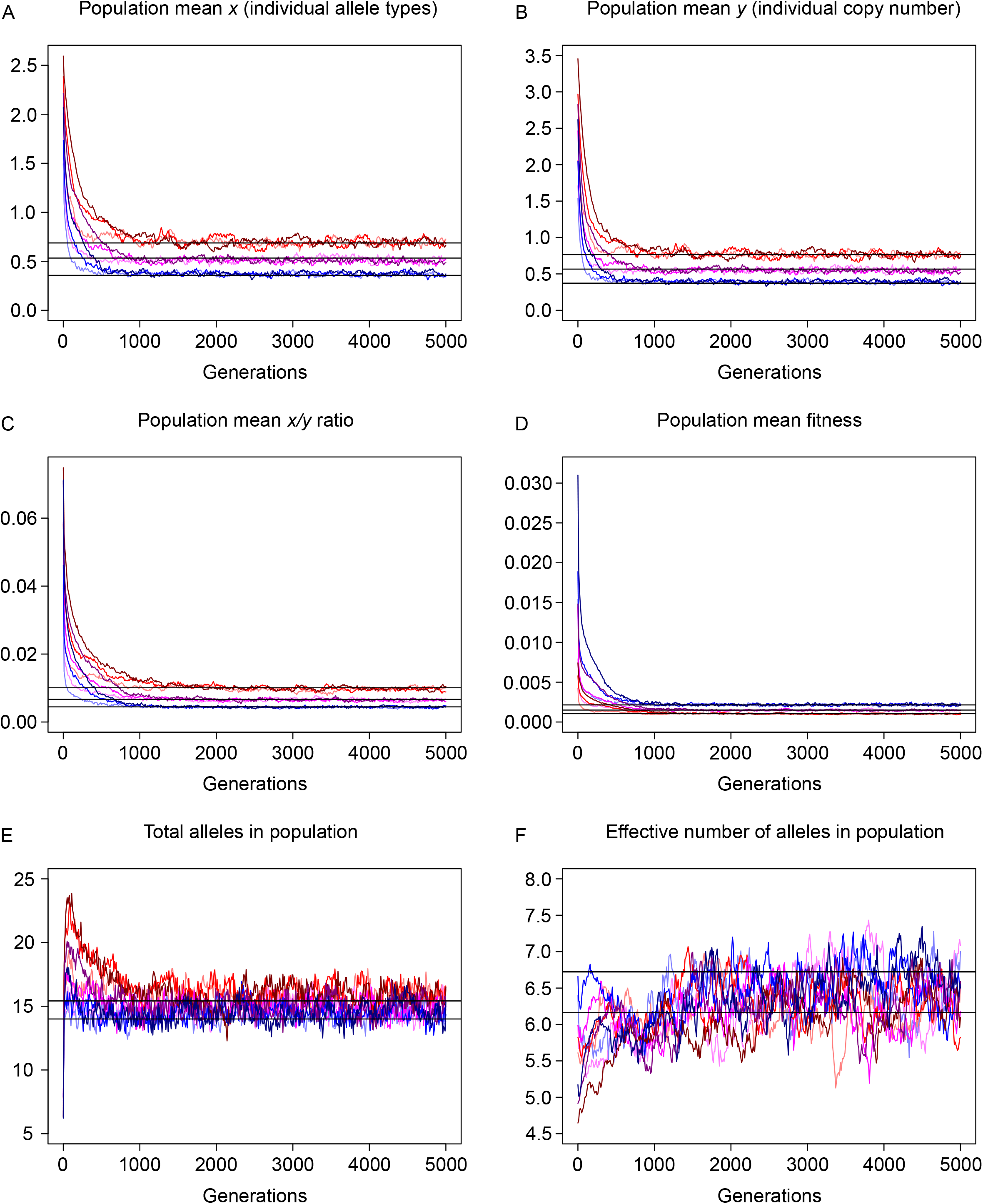
Standard deviation *among 200 replicates* from scenario (b) in six statistics during recovery from bottleneck, *N*_*e*_ = 2000. See Figure 6 for explanation of panels and colors.

**Figure S5.**
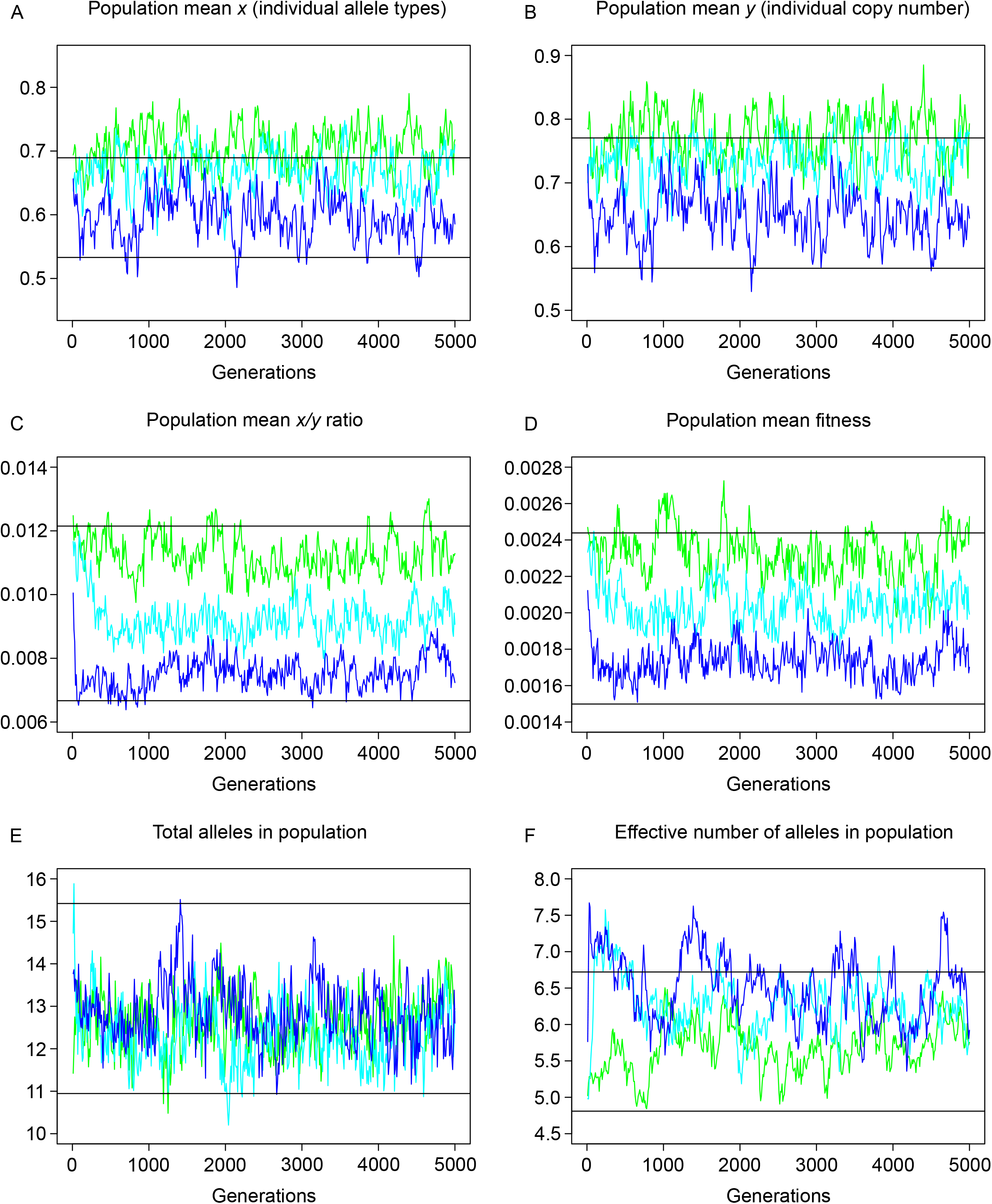
Standard deviation *among 2×100 replicates* from scenario (c) in six statistics after start of reciprocal migration, *N*_*e*_ *= 1000* in each subpopulation. See Figure 7 for explanation of panels and colors. Black lines indicate mean values (across 500 replicates) in panmictic populations of size *N*_*e*_ = 1000 (upper line for A-D, lower line for E-F) and *N*_*e*_ = 2000 (lower line for A-D, upper line for E-F).

**Figure S6.**
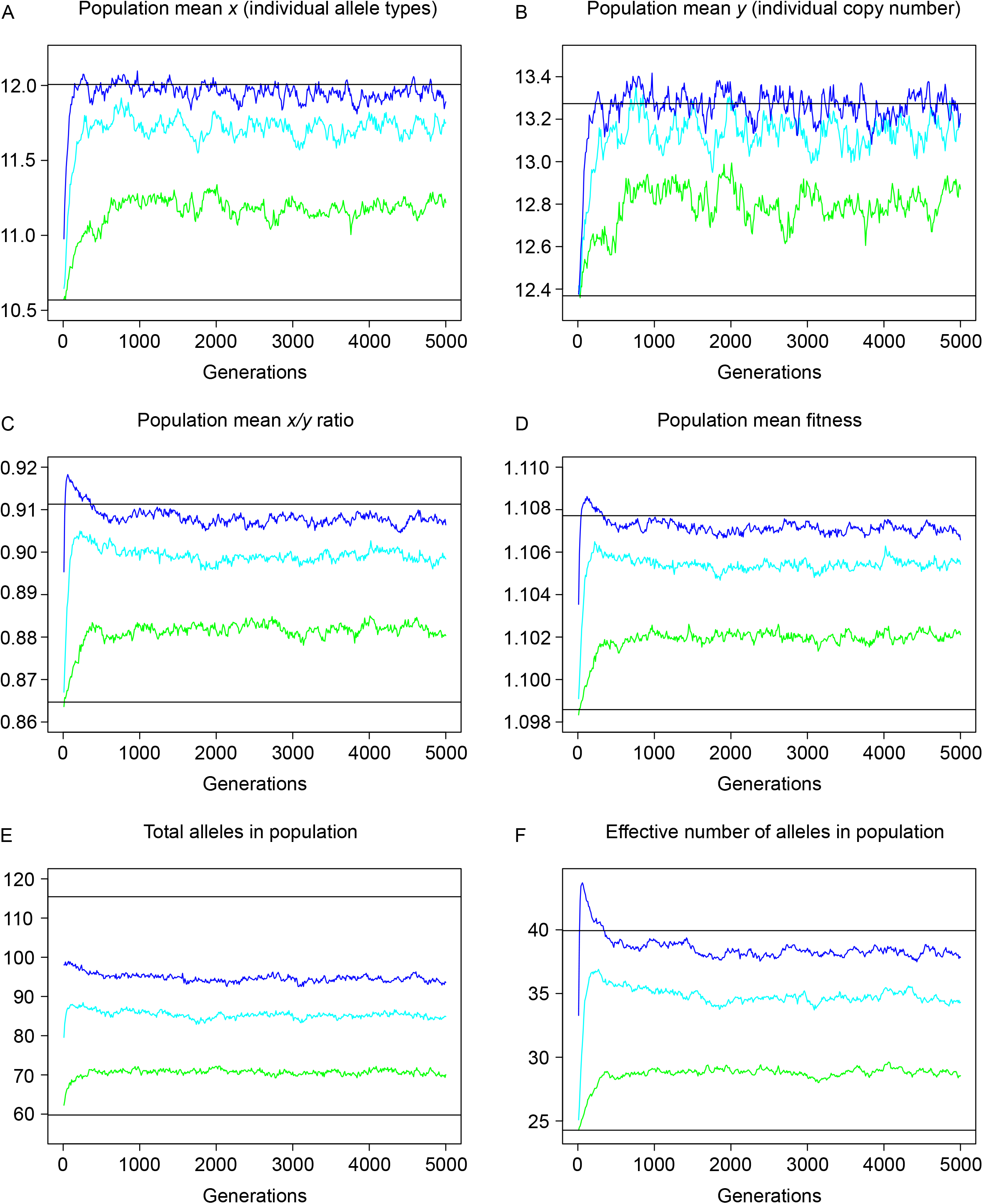
Mean values *across 2×100 replicates* from scenario (c) in six statistics after start of reciprocal migration, *N*_*e*_ *= 500* in each subpopulation. See Figure 7 for explanation of panels and colors. Black lines indicate mean values (across 500 replicates) in panmictic populations of size *N*_*e*_ = 500 (lower line) and *N*_*e*_ = 1000 (upper line).

**Figure S7.**
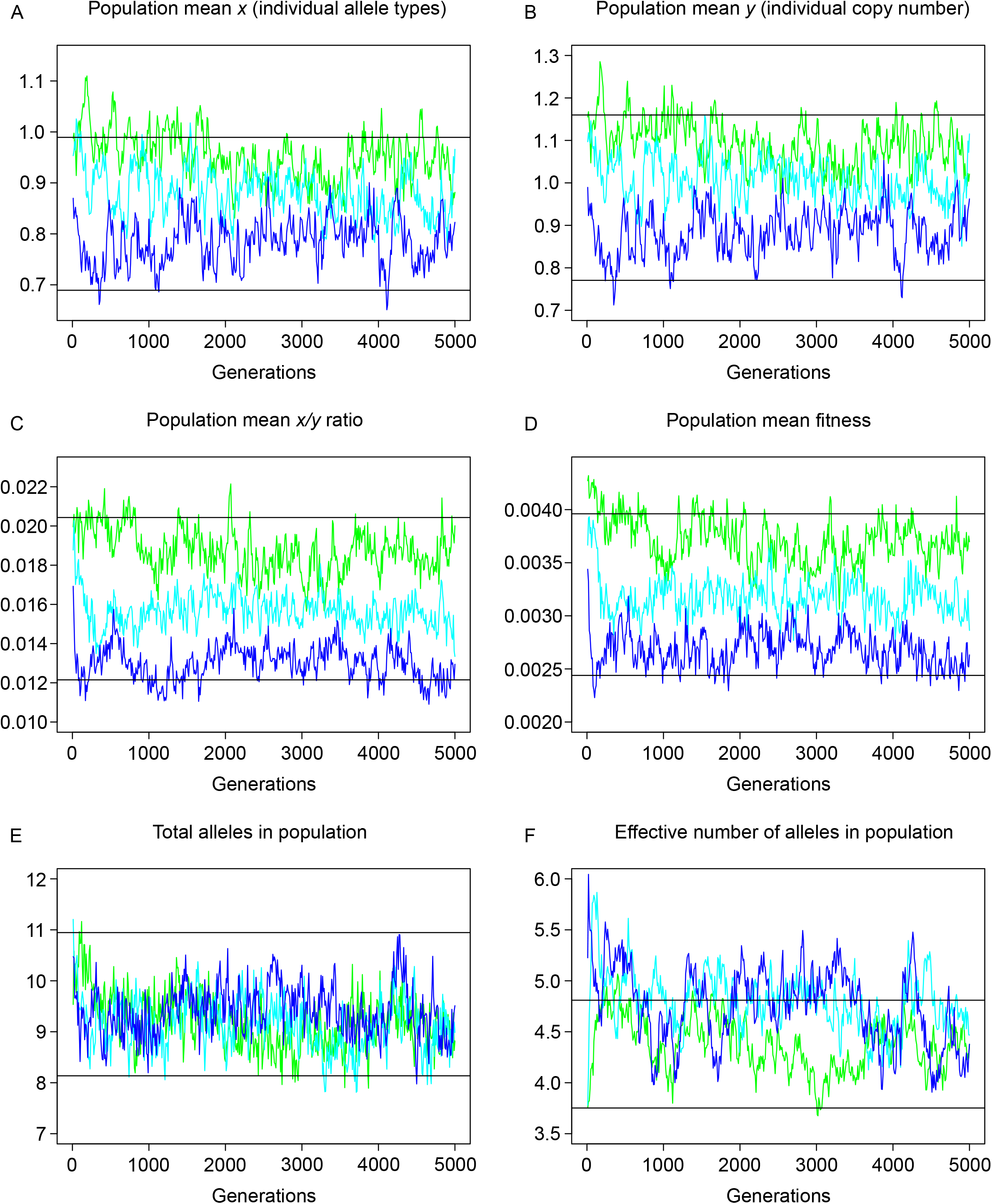
Standard deviation *among 2×100 replicates* from scenario (c) in six statistics after start of reciprocal migration, *N*_*e*_ *= 500* in each subpopulation. See Figure 7 for explanation of panels and colors. Black lines indicate mean values (across 500 replicates) in panmictic populations of size *N*_*e*_ = 500 (upper line for A-D, lower line for E-F) and *N*_*e*_ = 1000 (lower line for A-D, upper line for E-F).

